# The circadian clock is a pacemaker of the axonal regenerative ability

**DOI:** 10.1101/2023.08.07.552230

**Authors:** Francesco De Virgiliis, Franziska Mueller, Ilaria Palmisano, Jessica S Chadwick, Lucía Luengo Gutierrez, Angela Giarrizzo, Yuyang Yan, Matt Christopher Danzi, Carmen Picón Muñoz, Luming Zhou, Guiping Kong, Elisabeth Serger, Thomas Haynes Hutson, ines Maldonado, Yayue Song, Christoph Scheiermann, Marco Brancaccio, Simone Di Giovanni

## Abstract

Peripheral nervous system injuries lead to long-term neurological disability due to limited axonal regenerative ability. Injury-dependent and -independent physiological mechanisms have provided important molecular insight into neuronal regeneration. However, whether common molecular denominators underpin both injury-dependent and independent biological processes remain unclear. Here, we performed a comparative analysis of recently generated transcriptomic datasets associated with the regenerative ability of sciatic dorsal root ganglia (DRG). Surprisingly, circadian rhythms were identified as a the most significantly enriched biological process associated with regenerative capability. We demonstrate that DRG neurons possess an endogenous circadian clock with a 24h oscillations of circadian genes and that their regenerative ability displays a diurnal oscillation in a mouse model of sciatic nerve injury. Consistently, transcriptomic analysis of DRG neurons showed a significant time-of-day dependent enrichment for processes associated with axonal regeneration, development and growth, as well as circadian associated genes, including the core clock genes *Bmal1* and *Clock*. Indeed, DRG-specific ablation of the non-redundant clock gene *Bmal1* showed that it is required for regenerative gene expression, neuronal intrinsic circadian regeneration and target reinnervation. Lastly, lithium, a chrono-active compound, enhanced nerve regeneration, in wildtype but not in *Bmal1* and *Cry1/2*-deficient mice. Together, these data demonstrate that daily rhythms and the molecular clock fine-tune the regenerative response of DRG neurons, and that chrono-active drugs, such as lithium, are a novel potential approach to nerve repair.

## Introduction

Peripheral nervous system (PNS) injury is an increasingly common condition present across all ages, which is due to traumatic and non-traumatic causes. Axonal regeneration upon PNS injury is limited and lack of nerve re-connectivity typically results in permanent neurological disability^1,2^. Experimental evidence has shown that the regenerative ability is regulated by both injury-dependent mechanisms, such as the well-established conditioning lesion^3^ and the recently established enriched conditioning^4^ as well as physiological injury-independent mechanisms, such as environmental enrichment^5^ and dietary regimens^6,7^. While these studies led to the identification of molecular pathways affecting axonal regenerative ability, a unitary and comprehensive molecular understanding of axonal regeneration is lacking. Thus, we explored the intriguing possibility of a common molecular denominator across these different models and conditions, which could expand our fundamental understanding of axonal regeneration beyond the current knowledge. To this end, we performed an unbiased comparative analysis of enriched signalling pathways from our recently generated datasets associated with non-regenerative versus regenerative conditions. These include peripheral injuries, dietary regimens such as intermittent fasting^7^ and environmental enrichment^5^. Unexpectedly, we identified circadian rhythms as potential common biological mechanism regulating axonal regeneration. The circadian clock is a molecular time-keeping mechanism of ∼24h, based on transcription and translation feedback loops that drive clock gene expression and circadian outputs^8^. These include cycles of physiological and behavioural changes that anticipate daily, recurring occurrences of environmental stimuli^9^, such as light or food intake. More specifically, circadian rhythms regulate a vast array of physiological phenomena including transcription^10^, translation^11^, epigenetics^12^, inflammation^13,14^, metabolism^12,15^ as well as wound healing processes^16^, thus possessing the potential to regulate axonal regeneration. Despite the pervasive nature of circadian clocks, their role in axonal regeneration remains elusive. Here, we show that DRG neurons possess an intrinsic molecular clock machinery that is necessary to time and optimize axonal regeneration and target re-innervation in a mouse model of sciatic nerve injury. Mechanistically, circadian axonal regeneration requires the neuronal expression of the core clock protein Bmal1, which regulates the regenerative program of sensory neurons.

Finally, we identified the potential of promoting axonal regeneration after injury by a clock-dependent mechanism, by using the chrono-active drug lithium, currently approved for treating neurological disorders^17–19^.

Together, our findings indicate that circadian rhythms tune the axonal regenerative ability to specific time windows of the day. This new knowledge could pave the way for the design of chrono-active regenerative therapies and neurorehabilitation regimens.

## Results

### Time-of-day dependent axon regeneration after sciatic nerve injury

We first aimed to determine the presence of a potential common molecular denominator across different regeneration models and conditions affecting the DRG. To this end, we performed a comparative transcriptomic analysis taking advantage of previously generated RNA sequencing datasets from established non-regenerative spinal dorsal column axotomy (DCA) compared to regenerative models^4,5,20^. Specifically, we included the regenerative injury-dependent conditioning lesion (sciatic nerve axotomy (SNA^20^) and sciatic nerve crush (SNC)). We further incorporated environmental enrichment and SNA (EE+SNA^4^, also known as enriched conditioning), as well as regenerative injury-independent models such as environmental enrichment (EE^5^) and intermittent fasting (IF)^7^. We investigated gene ontology (GO) functional categories and ranked them by their enrichment in regenerative versus the DCA non-regenerative datasets (**Figure 1A**). Unexpectedly, comparative GO analyses (Fisher test, P-value < 0.05) of these datasets showed that “Circadian rhythms” was the only enriched biological process shared across all the regenerative datasets analysed that was not enriched in non-regenerative DCA (**Figure 1A**). Therefore, we hypothesised that the circadian clock may influence the regenerative capacity of DRG neurons depending on the time of the day at which the injury is performed. First, we assessed whether daily variations of clock gene expression could be detected in unchallenged DRG by monitoring the expression levels of the core clock genes *Bmal1* (encoded by *Arntl*), *Clock*, *Per1*, *Per2*, *Cry1* and *Cry2* across the day in naïve DRG. We performed experiments every 4h at *Zeitgeber* times (ZTs) 0, 4, 8, 12, 16, 20 and 24 (i.e. hours after light onset in a 12h light / 12h dark environment), corresponding to 7am, 11am, 3pm, 7pm, 11pm and 3am), and found clock gene expression levels display diurnal oscillations (JTK_CYCLE, p<0.05 for all genes; **Figure 1B**, **Table 1**), in line with previous reports ^21^. Next, we assessed whether the DRG response to injury would differ depending on the time of the day at which the injury is performed. We performed a SNC injury at ZT 0, 4, 8, 12, 16, 20 and 24 and assessed sciatic nerve regeneration 24 hours and 72 hours post injury by measuring sciatic regeneration index and fluorescence intensity along the nerve of the regenerative marker SGC10 (**Figure 1C**). Importantly, we observed a significant increase in SCG10 positive regenerating fibres and sciatic regeneration index when the injury was performed at ZT20, compared to other time points, both at 24 hours (**Figure 1D-E and Supplementary Figure 1A-B**) and 72 hours post-injury (**Figure 1F-G and Supplementary Figure 1C-D**). Given the high heterogeneity of sensory fibres that compose the sciatic nerve, we asked whether different neuronal subsets in the DRG would account for the phenotype we observed. For this, we injected the retrograde tracer cholera toxin B (CTB) in the sciatic nerve where we find the fibres originating in the DRG. Given CTB is injected distally to the lesion, only regenerating fibres are capable of uptaking the tracer, allowing for direct and unequivocal measurement of regenerating DRG neurons. Subsequently, DRG neurons were immunostained with neurofilament-200 (NF-200), as a marker of large diameter proprioceptive neurons as well as calcitonin gene-related peptide (CGRP) as marker of small diameter nociceptors. In line with our previous data, we found that the number of CTB+ DRG neurons (i.e. regenerating neurons) was significantly increased at ZT20 compared to ZT8 (**Supplementary Figure 2A-B**), indicating higher regeneration potential at ZT20. However, the distribution of CTB labelled DRG neurons in the two neuronal subsets remained the same across the conditions (**Supplementary Figure 2A-C**), suggesting that time-of-day differences in the regenerative response affects proprioceptive and nociceptive populations equally.

**Figure 1.**
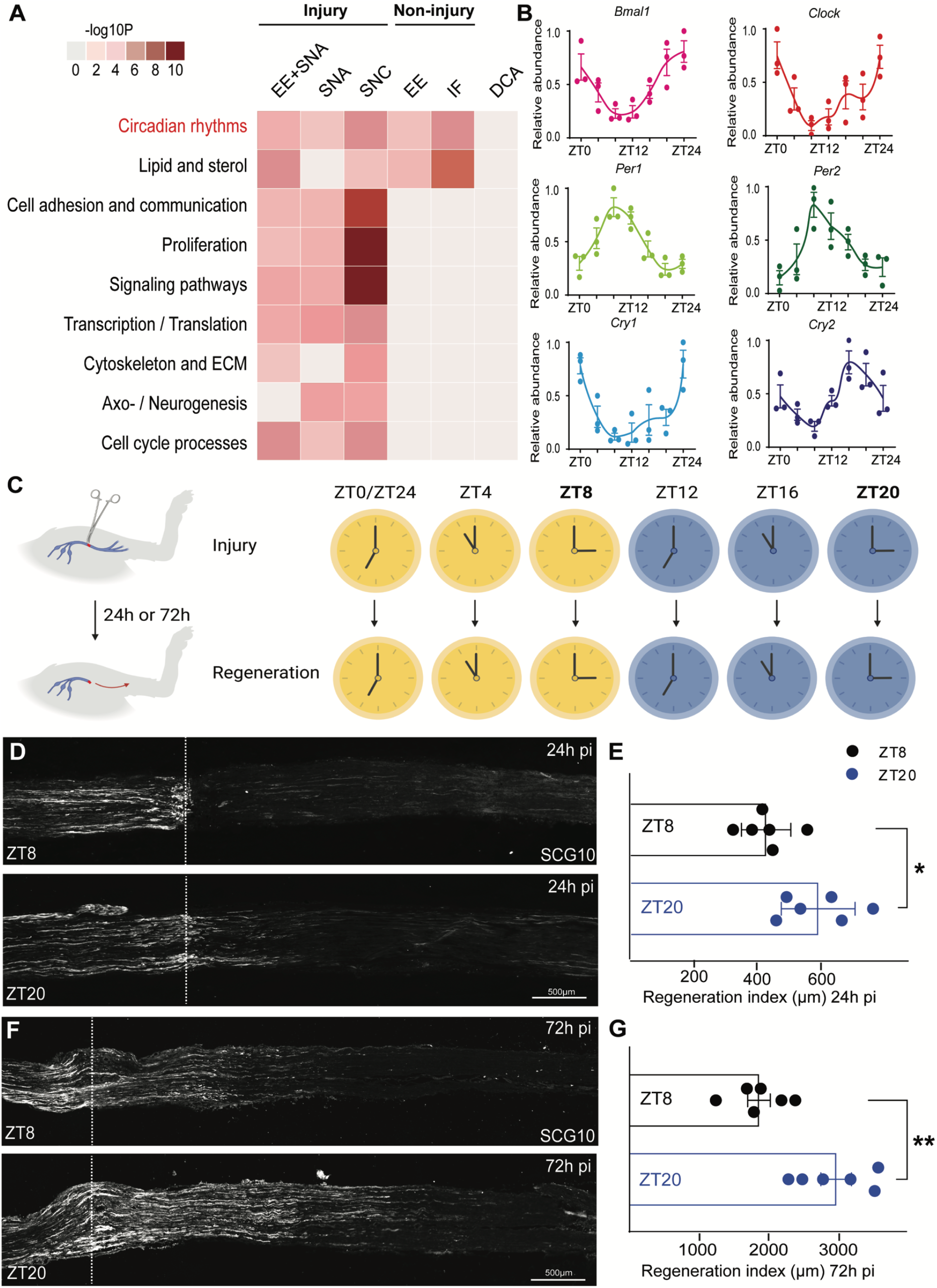
Circadian rhythms are common regulators of axonal regeneration. **A.** Heatmap showing the Gene Ontology analysis (molecular function, DAVID) of the DE genes modulated by injury-dependent, injury-independent regenerative and non-regenerative (DCA) conditions (modified Fisher’s exact, p<0.05), (EE=Enriched environment, SNA=sciatic nerve axotomy, SNC=Sciatic nerve crush, IF=intermittent fasting, DCA=dorsal column axotomy). **B.** Quantitative RT-PCR analysis of the mRNA levels of the circadian clock genes *Bmal1*, *CLOCK*, *Per1*, *Per2*, *Cry1* and *Cry2* mRNA levels in DRG at ZT0, ZT4, ZT8, ZT12, ZT16, ZT20 and ZT24 and normalised over GAPDH (mean ± SEM, JTK_CYCLE analysis, p<0.05, n=3 biologically independent animals/group examined over 3 independent experiments). **C.** Timeline for the *in vivo* experiment. **D.** Representative images of sciatic nerves injured at ZT8 or ZT20 stained for SCG10 24h after SNC (pi=post injury). Scale bar, 500 µm. **E.** Quantification of SCG10 intensity from the lesion site shown as regeneration index (distance from the injury site (dotted lines) at which 50% of the fluorescence decays) (one-way ANOVA, Tukey’s post-hoc, p<0.05, n = 6 biologically independent animals/group). Fluorescence intensity was measured in one series of tissue sections for each nerve. **F.** Timeline for the *in vivo* experiment. **G.** Representative images of sciatic nerves injured at ZT8 or ZT20 stained for SCG10 72h after SNC (pi=post injury). Scale bar, 500 µm. **H.** Quantification of SCG10 intensity from the lesion site shown as regeneration index (distance from the injury site (dotted lines) at which 50% of the fluorescence decays) (one-way ANOVA, Tukey’s post-hoc, p<0.05, n = 6 biologically independent animals/group). Fluorescence intensity was measured in one series of tissue sections for each nerve.

**Table 1.**
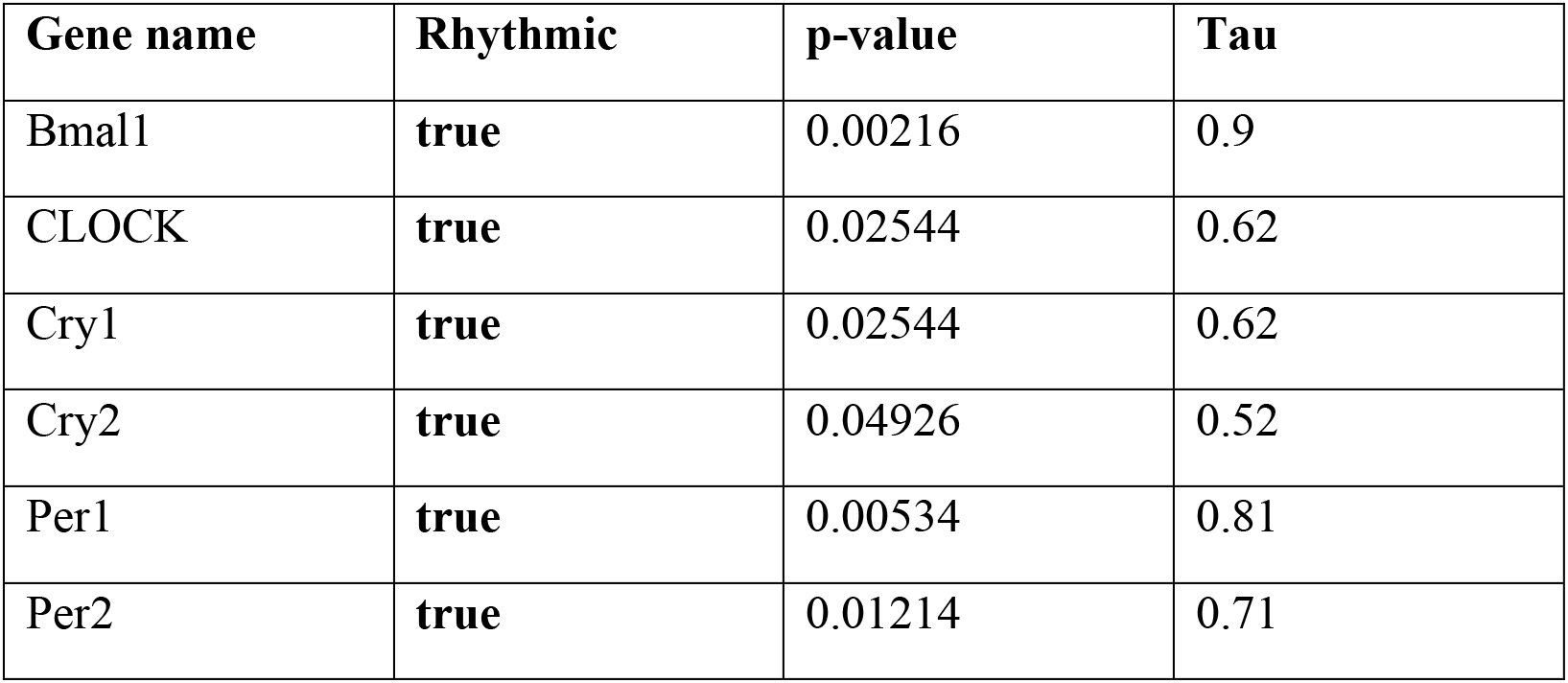
Analysis of rhythmicity of clock genes qPCR data represented in Figure 1B. JTK_CYCLE on mean values, with dataset compared to a 24h cosine wave with a 4h phase spread, p-value<0.05.

| Gene name | Rhythmic | p-value | Tau |
| --- | --- | --- | --- |
| Bmal1 | true | 0.00216 | 0.9 |
| CLOCK | true | 0.02544 | 0.62 |
| Cry1 | true | 0.02544 | 0.62 |
| Cry2 | true | 0.04926 | 0.52 |
| Per1 | true | 0.00534 | 0.81 |
| Per2 | true | 0.01214 | 0.71 |

Studies in neuronal and non-neuronal tissues showed that diurnal oscillations of clock genes modulate cells and organ responses, including inflammation, cell proliferation, hormonal levels as well as injury responses^9,13,14,16^. Thus, we asked whether non-neuronal cells including macrophages and neutrophils as well as nerve-associated Schwann Cells (SCs), known to play a role in sciatic nerve regeneration after injury ^22–24^, were modulated by the time-of-day at which the injury was performed. We found that macrophage (CD68^+^) and neutrophil (Ly6G^+^) infiltration at the injury site 24h or 72h after injury was unchanged (**Supplementary Figure 3A-H and Supplementary Figure 5**). Likewise, we did not observe time-of-day-dependent changes in SCs (SOX10^+^) proliferation 72h after injury, when SCs typically proliferate (**Supplementary Figure 4A-B and Supplementary Figure 5**). Additionally, since neurotrophins are important molecules involved in the regenerative response of DRG neurons after peripheral nerve injury^25,26^, we measured the levels of the neurotrophins NT3, NT4/5, BDNF and NGF in naïve DRG harvested at the times of trough (ZT8) and peak (ZT20) regeneration potential and observed no changes (**Supplementary Figure 6**).

Together, these data show that the regenerative ability of DRG neurons displays a time-of-day dependent regulation and suggest the presence of a neuronal intrinsic endogenous clock that governs the injury response of DRG neurons.

### Transcriptomic analysis of injured DRG neurons shows time-dependent enrichment for circadian and developmental growth biological processes associated with high regeneration potential

Next, we investigated the time-of-day dependent transcriptional landscape of DRG neurons associated with the peak (ZT20) and trough (ZT8) in regenerative ability, which positively correlates with Bmal1 expression **(Supplementary Figure 7).** To this end, we carried out RNA-sequencing (RNA-seq) 72h after a nerve injury was performed at the times of regenerative trough (ZT8) or peak (ZT20) (**Figure 2A**). To specifically enrich for neuronal transcripts, the DRG suspension was selectively enriched for neurons using a BSA gradient (**Figure 2A**). Principal component analysis (PCA) and unsupervised clustering showed a clear separation between the two experimental groups ZT8 vs ZT20 (**Figure 2B and Supplementary Figure 8A**). Remarkably, gene ontology (GO) analysis showed a significant enrichment for biological processes associated with chromatin organization and transcription, axonogenesis and axon guidance, neuronal development, cytoskeleton, synaptic regulation and cell adhesion (**Figure 2C and Supplementary File 1 and 2**). Circadian rhythms and clock genes were also enriched, with differentially expressed clock genes including *Arntl* (*Bmal1)*, *Clock*, *Timeless*, and *Bhlhe40/41* (upregulated), as well as *Nr1d1*, *Usp1*, and *Dbp* (downregulated) (**Figure 2D, Supplementary File 1 and Table 2**), supporting the robustness of the dataset and suggesting a possible role for the molecular clock in regulating nerve regeneration. Furthermore, developmental growth genes, such as *Notch2/3*, *Ephrin a4/5* and *Robo1/2* family members as well as classical regeneration associated genes (RAGs) were significantly upregulated at ZT20 vs ZT8. They include the transcription factors c-*myc*, several *Klf* family members, *Atf5*, *Stat3*, *Creb1*, *Creb1-bp* (CBP), *Hif1a*, *FOXO3a* and P53 target genes *Tp53inp1/2*; the cytoskeletal and membrane associated factors *Basp1*, *Ncam1*, *Coronin2a*, *Map1a* and *Map2*; the growth factors *Ngf-rec*, *Vgf*, T*gf-rec, Hgf*, *Fgf11*, *Fgf14* and *Egf*; the signalling molecules *Jak1*, *Prkce* and *Prkcg*, as well as molecules belonging toIL6 signalling such as *Il6st,* among others. (**Figure 2D, Supplementary File 1 and Table 2**). Additional downregulated RAGs, whose inhibition promotes axon regeneration, include the histone deacetylases *Hdac1* and *Hdac2*, *Cacna1s* as well as the GSK3-β signalling associated molecule *Gskip* (**Figure 2D, Supplementary File 1 and Table 2**).

**Figure 2.**
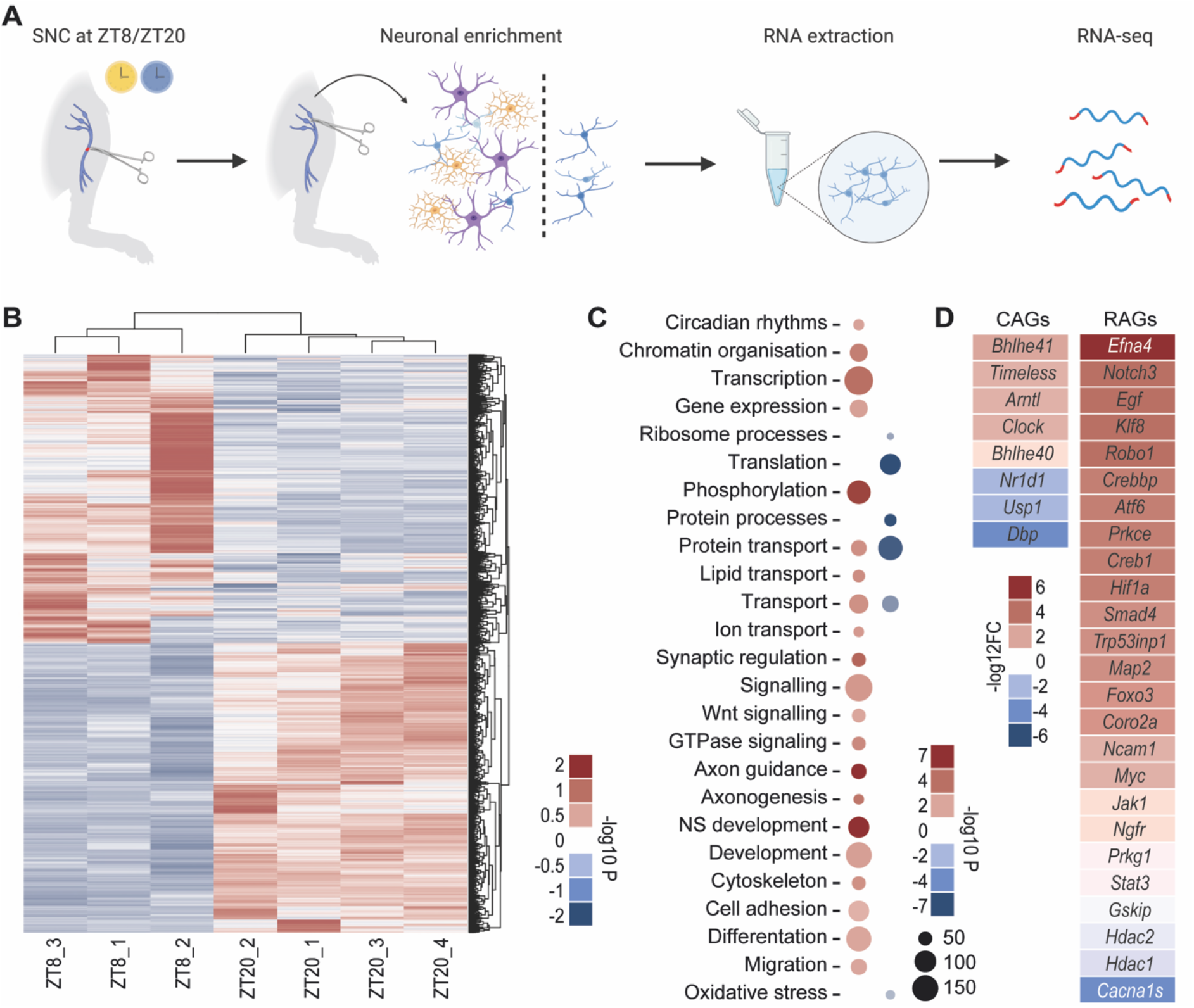
Transcriptional analysis in DRG after injury identifies time-of-day dependent processes associated with axonal regeneration. **A.** Timeline for the transcriptomic experiment. **B.** Heatmap of differentially expressed genes (FDR<0.01) belonging to DRG neurons injured at ZT20 vs ZT8, analysed 72h after injury. **B.** Dendrogram of unsupervised clustered DE genes across all experimental conditions (ZT8 vs ZT20, -log10 adjusted p-value<0.05, red=upregulated, blue=downregulated). **C.** Dot-plot showing the Gene Ontology analysis (biological processes, DAVID) of the DE genes modulated by a ZT8 or ZT20 nerve injury in DRG neurons (-log10 adjusted p-value<0.05, red=upregulated, blue=downregulated). **D.** Expression levels of Circadian associated genes (CAGs) and regeneration associated genes (RAGs) (-log2 Fold Change, red=upregulated, blue=downregulated) (EE=Enriched environment, SNA=sciatic nerve axotomy, SNC=Sciatic nerve crush, IF=intermittent fasting, DCA=dorsal column axotomy).

**Table 2.** Extended table reporting gene expression data (gene name, fold change, p-value) of DE Clock associated genes (CAGs) and Regeneration associated genes (RAGs) found in the RNA-seq of DRG after an injury performed at ZT20 vs ZT8 (see also **Figure 2)**.

| CAGs | Fold Change | p-value |
| --- | --- | --- |
| Bhlhe41 | 1.801257 | 2.30E-05 |
| Timeless | 0.91587 | 0.02544 |
| Arntl | 0.812613577 | 0.02544 |
| Clock | 0.801852 | 0.04926 |
| Bhlhe40 | 0.609778679 | 0.00534 |
| Nr1d1 | -1.273298213 | 0.01214 |
| Usp1 | -1.26362 | 0.047242 |
| Dbp | -2.70571 | 2.08E-05 |
| RAGs | Fold Change | p-value |
| Efna4 | 5.792264 | 0.019385 |
| Notch3 | 2.568564 | 0.012489 |
| Egf | 2.230509 | 0.029209 |
| Klf8 | 2.22704 | 0.023871 |
| Fgf14 | 2.148082 | 6.60E-12 |
| Notch2 | 2.028718 | 1.18E-05 |
| Klf1 | 1.837033 | 0.017693 |
| Robo1 | 1.534673 | 0.000704 |
| Klf13 | 1.527326 | 4.67E-05 |
| Crebl2 | 1.105803 | 0.016278 |
| Efnb2 | 1.043021 | 5.72E-07 |
| Crebbp | 1.031487 | 0.000416 |
| Fgf11 | 1.000311 | 0.012039 |
| Robo2 | 0.970628 | 5.21E-05 |
| Klf5 | 0.93096 | 0.037397 |
| Atf6 | 0.929285 | 0.002073 |
| Klf16 | 0.924909 | 0.000936 |
| Prkce | 0.916592 | 0.001676 |
| Creb1 | 0.903214 | 1.39E-09 |
| Hif1a | 0.890155 | 0.000877 |
| Hgf | 0.868469 | 0.036293 |
| Efna5 | 0.853414 | 0.01086 |
| Smad4 | 0.837186 | 0.003453 |
| Trp53inp1 | 0.821017 | 0.000381 |
| Map2 | 0.812847 | 0.000776 |
| Basp1 | 0.792412 | 0.009105 |
| Vgf | 0.77967 | 0.039477 |
| Atf5 | 0.7454 | 0.000188 |
| Foxo3 | 0.708334 | 0.019678 |
| Klf12 | 0.696557 | 0.019584 |
| Il6st | 0.686253 | 0.008956 |
| Fosl2 | 0.680117 | 0.005823 |
| Trp53inp2 | 0.670738 | 0.009432 |
| Map1a | 0.661544 | 0.03181 |
| Tgfbr1 | 0.631607 | 0.039091 |
| Coro2a | 0.618138 | 0.001622 |
| Ncam1 | 0.594811 | 0.011897 |
| Myc | 0.560053 | 0.002388 |
| Ntrk3 | 0.540458 | 0.001776 |
| Prkcd | 0.514291 | 0.005399 |
| Jak1 | 0.501427 | 0.000182 |
| Itgb1 | 0.482534 | 0.000292 |
| Ngfr | 0.465999 | 0.023534 |
| Prkg1 | 0.449758 | 0.044489 |
| Trp53bp1 | 0.443054 | 0.023916 |
| Klf7 | 0.425923 | 0.000199 |
| Stat3 | 0.349548 | 0.024309 |
| Gskip | -0.3778 | 0.002197 |
| Hdac2 | -0.54748 | 0.034359 |
| Hdac1 | -0.76848 | 1.08E-05 |
| Cacna1s | -2.70701 | 0.023011 |

Importantly, odds ratio analysis, which measures the statistical correlation between datasets, showed that this RNA-seq dataset correlates significantly with RNA-seq datasets associated with regenerative states (SNA, SNC, IF), but not with a non-regenerative state (DCA) (**Supplementary Figure 8B**). Together, these data show a time-of-day change in transcriptional landscape of DRG neurons after injury and support a role for the molecular clock in regulating the regenerative ability of DRG neurons.

### A Bmal1 dependent clock regulates the regenerative growth of DRG neurons

Based on the RNAseq dataset, we asked whether DRG neurons possess an endogenous oscillating clock and whether we can perturb its function via genetic deletion of the non-redundant core clock protein BMAL1^27^. To this aim, we took advantage of Period 2-Luciferase (Per2-Luc) mice where Per2 expression is used as proxy of clock gene oscillation and is quantified by a co-expressed Luciferase reporter. To functionally disrupt the circadian clock *in vitro*, dissociated neuronal enriched DRG were transfected with siRNA anti-*Bmal1* to knock down (KD) BMAL1 protein expression (**Supplementary Figure 9**), or scrambled control. 48h after transfection, DRG cultures were synchronised by a 2h 100nM Dexamethasone (Dexa) pulse as previously reported^28,29^. 24 hours after synchronization (T=24), luciferase luminescence in neuronal enriched cultures was assessed every 4 hours across a 24-hour period (**Figure 3A**). We observed a rhythmic oscillation in Per2-Luciferase expression in control cultures, which was abolished in BMAL1 KD cultures (JTK_CYCLE, p<0.005) (**Figure 3 B-C and Table 3**), demonstrating the presence of an endogenous BMAL1-dependent clock in DRG neurons.

**Figure 3.**
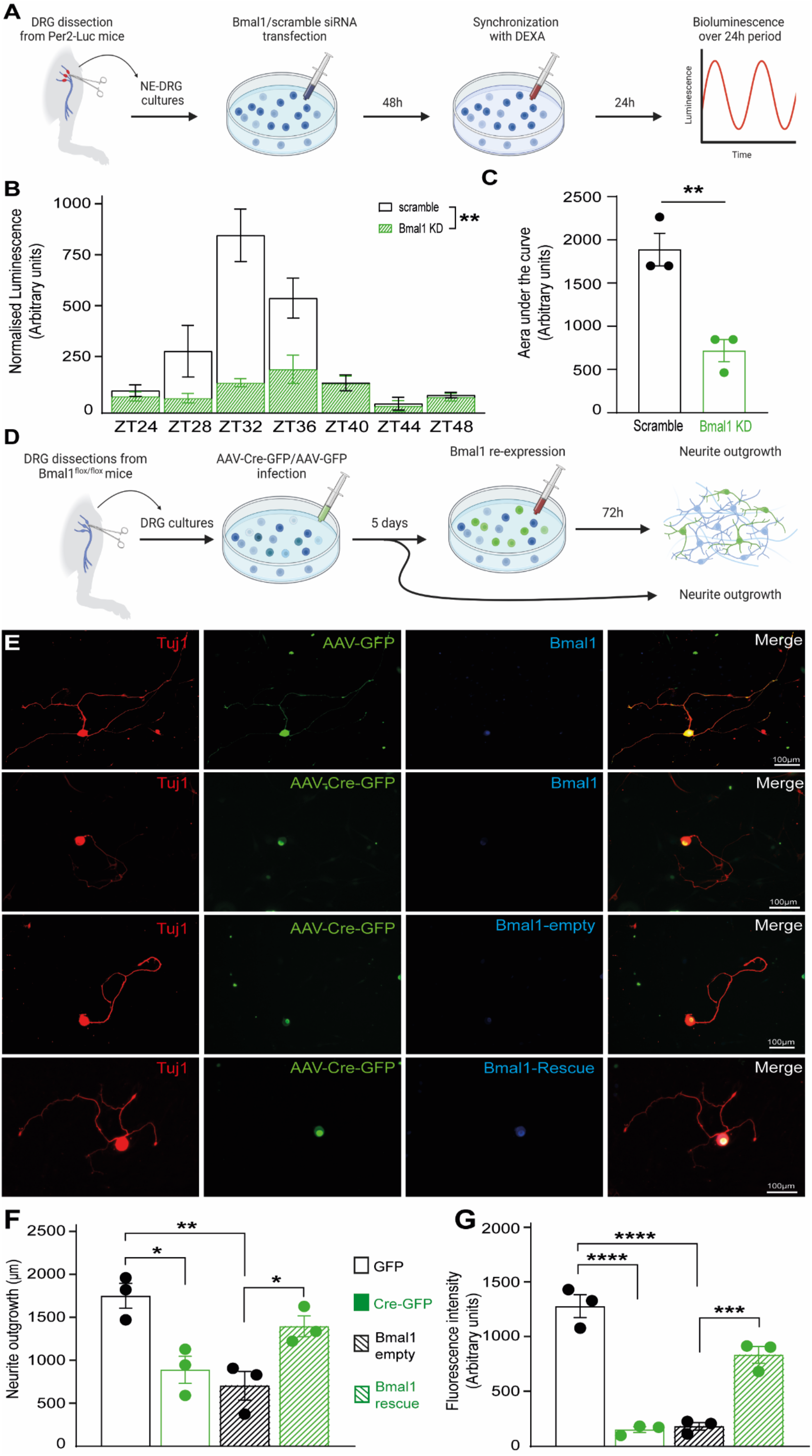
Bmal1 controls the molecular clock and regenerative growth of DRG neurons. **A.** Timeline for the *in vitro* experiment. **B.** Representation of normalised luminescence in DRG neuronal cell cultures from Per-Luc animals transduced with siRNA anti-*Bmal1* vs scramble and analysed at ZT0, ZT4, ZT8, ZT12, ZT16, ZT20 (JTK CYCLE analysis, p<0.005, n = 3 biologically independent experiments /group). **C.** Histogram quantification of the area under the curve of the experiment in B (Student T-test, p<0.05). **D.** Timeline for the *in vitro* experiment. **E.** Representative images of cultured DRG neurons from *Bmal1*^flox/flox^ infected with GFP vs Cre-GFP and immunostained for Tuj1 (red), GFP (green) and BMAL1 (blue). Scale bar, 100 µm. **F.** Quantification of neurite outgrowth shown as neurite outgrowth in μm (one-way ANOVA, Tukey’s post-hoc, p<0.05, n = 3 biologically independent experiments/group). **G.** Quantification of Bmal1 fluorescence (one-way ANOVA, Tukey’s post-hoc, p<0.05, n = 3 biologically independent experiments /group).

**Table 3.** Analysis of rhythmicity of Per2-Luciferase data represented in Figure 3B. JTK_CYCLE on mean values, with dataset compared to a 24h cosine wave with a 4h phase spread, p-value<0.005.

| Group | Rhythmic | p-value | Tau |
| --- | --- | --- | --- |
| Scramble 1 | false | 0.0414 | 0.55 |
| Scramble 2 | true | 0.00367 | 0.85 |
| Scramble 3 | true | 0.0028 | 0.88 |
| Bmal1 KD 1 | false | 0.0574 | 0.5 |
| Bmal1 KD 2 | false | 0.0414 | 0.55 |
| Bmal1 KD 3 | false | 0.06191 | 0.49 |
| Scramble mean | true | 0.0028 | 0.88 |
| Bmal1 KD mean | false | 0.06191 | 0.49 |

In order to selectively analyse neurons where BMAL1 was knocked down *in vitro*, we transfected dissociated neuronal enriched DRG cultures from *Bmal1*^flox/flox^ mice with an AAV-Cre-GFP or control AAV-GFP for 5 days before re-plating them (**Figure 3D**). After 24 hours in culture, we observed that Cre-GFP infected DRG (*Bmal1* cKO) neurons displayed significantly attenuated neurite outgrowth compared to GFP controls (**Figure 3E-F**). Additionally, Bmal1 expression was significantly reduced in Cre-GFP neurons compared to GFP controls (**Figure 3E-G**). When we re-induced BMAL1 expression in DRG neurons by transfecting AAV-Cre-GFP infected cultured DRG with a *Bmal1* over-expression plasmid (**Figure 3D**), we observed a significant rescue of the neurite outgrowth phenotype correlated with a rescued BMAL1 protein expression. (**Figure 3E-G**). Importantly, the number of neurons did not differ between experimental conditions indicating equal survival rates **(Supplementary Figure 10).** Together, these data demonstrate that BMAL1 controls the endogenous clock and the regenerative growth of DRG sensory neurons.

### Bmal1 deletion abolishes circadian-dependent axon regeneration and target re-innervation of DRG neurons

We next investigated whether a functional clock was required for the time-of-day dependent DRG regeneration *in vivo*. To functionally disrupt the molecular clock *in vivo*, we conditionally deleted *Bmal1* in DRG neurons by injecting an AAV-Cre-GFP or control AAV-GFP in the sciatic nerve of Bmal1^flox/flox^ mice (cKO) 4 weeks before performing a SNC at the regeneration peak (ZT20) and trough (ZT8) (**Supplementary Figure 11 and Supplementary Figure 12**), and assessed sciatic nerve regeneration 72 hours after injury (**Figure 4A**). Importantly, similarly to in cell culture data, *Bmal1* deletion did not lead to neuronal toxicity, given that the number of neurons and apoptotic nuclei did not differ between conditions (**Supplementary Figure 13).** Additionally, to directly assess regeneration of infected DRG neurons, CTB was injected distally to the lesion site at the time of the injury and CTB positive GFP transduced DRG neurons were analysed, allowing for direct and unequivocal measurement of regenerating DRG neurons. While control virus injected animals reproduced the time-of-day dependent phenotype in neuron regeneration, this was completely abrogated in Cre-virus-injected cKO animals, with the ZT20 regeneration capacity being reduced to ZT8 levels (**Figure 4B-D and Supplementary Figure 11**). To assess whether *Bmal1* deletion in DRG neurons affects sensorimotor behaviour, which might explain the phenotype observed, we quantified locomotor activity using an open field test and sensorimotor function with a grid walk test prior to injury at ZT8 and ZT20 in wild-type (WT) and *Bmal1* cKO animals. We observed increased locomotor activity when mice were tested during their active phase (ZT20) compared to their resting phase (ZT8) (**Supplementary Figure 14B**). However, locomotor activity and sensorimotor function were unaltered in *Bmal1* cKO compared to WT animals (**Supplementary Figure 14A-B**). These data indicate that the BMAL1-dependent time-of-day differences in regeneration are not due to *Bmal1* cKO behavioral alterations. Additionally, we investigated whether *Bmal1* was required for the expression of classical RAGs^30,31^ whose expression is induced by a nerve injury versus sham surgery and whose manipulation has previously been found to affect the regenerative ability of DRG neurons^30,31^. In line with the nerve regeneration and RNA-seq data, we found that expression of *c-myc*, *Creb*, *Stat3*, *Tp53* and *Il6* signalling was also induced at ZT20 vs ZT8 (**Supplementary Figure 15**). In addition, we observed reduced expression levels in *Atf3*, *c-jun*, *c-myc*, *c-fos*, *Creb*, *Stat3* in *Bmal1* cKO when SNC was performed at ZT20, while no changes were observed in *Tp53*, *Il6*, *Rac1* or the control gene *β3-tubulin* expression levels (**Supplementary Figure 15C**). By immunostaining, we observed that the levels of pCREB and pS6 were increased when an injury was performed at ZT20 compared to ZT8 and that this increase was abolished in *Bmal1* cKO DRG compared to controls (**Supplementary Figure 15**). The regeneration-associated epigenetic mark (RAE) H3K27ac^32^, which is acetylated by CBP (upregulated in ZT20 vs ZT8 in DRG neurons after injury), was also increased at ZT20 vs ZT8 in WT but not in *Bmal1* cKO DRG compared to controls (**Supplementary Figure 16**). Together, these data suggest that *Bmal1* regulates the regenerative transcriptional response of DRG neurons.

**Figure 4.**
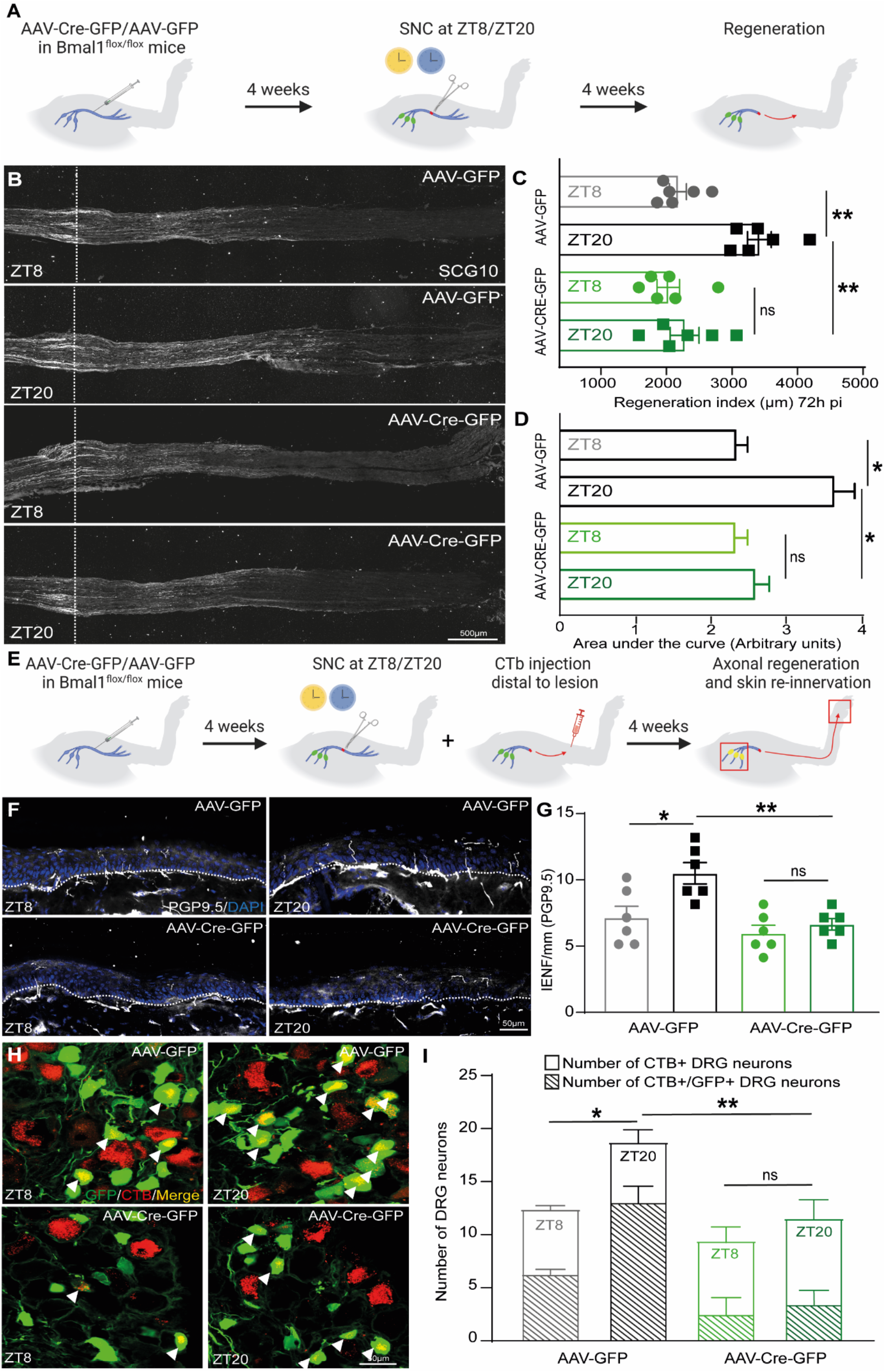
Bmal1 controls the regenerative ability and target re-innervation of DRG neurons *in vivo*. **A.** Timeline for the *in vivo* experiment. **B.** Representative images of sciatic nerves from Bmal1^fl/fl^ animals infected with GFP vs Cre-GFP virus injured at ZT8 or ZT20 immunostained for SCG10 72h after SNC. Scale bar, 500 µm. **C.** Quantification of SCG10 intensity from the lesion site shown as regeneration index (distance from the injury site at which 50% of the fluorescence decays) (two-way ANOVA, Tukey’s post-hoc, p<0.05, n = 4 biologically independent animals/group). Fluorescence intensity was measured in one series of tissue sections for each nerve. **D.** Quantification of the SCG10 fluorescence along the nerves starting from the lesion sites (dotted lines) visualised as area under the regeneration curve (two-way ANOVA, Tukey’s post-hoc, p<0.05, n = 6 biologically independent animals/group). Fluorescence intensity was measured in one series of tissue sections for each nerve. **E.** Timeline for the *in vivo* experiment. **F.** Representative images of skin sections from Bmal1 animals infected with GFP vs Cre-GFP virus injured at ZT8 or ZT20 stained for PGP9.5 21days after SNC. Scale bar, 50 µm. **G.** Quantification of PGP9.5 positive nerve endings in the epidermis (above dotted lines) (two-way ANOVA, Tukey’s post-hoc, p<0.05, n = 4 biologically independent animals/group). Number of fibres was measured in one series of tissue sections for each skin section. **H.** Representative images of DRGs from Bmal1 animals infected with PGFP vs Cre-GFP (green) virus injured at ZT8 or ZT20, injected with CTB retrograde tracer (red) and analysed 21days after SNC. Arrowheads highlight double positive DRG neurons (yellow). Scale bar, 50 µm. **I.** Quantification of CTB and GFP double positive DRG neurons (two-way ANOVA, Tukey’s post-hoc, p<0.05, n = 6 biologically independent animals/group). Number of cells was measured in one series of tissue sections for each DRG.

While injured adult PNS fibres can mount a regenerative response, nerve regeneration is slow and often incomplete, resulting in lack of target reinnervation and long-term disability^1,2,25^. Thus, we asked whether the regenerative gains observed up to 72 hours after SNC would result in improved long-term target re-innervation and whether this effect was due to the presence of a functional *Bmal1*-dependent clock in DRG neurons. Therefore, we conditionally deleted Bmal1 in DRG neurons by injecting an AAV-Cre-GFP or control AAV-GFP in the sciatic nerve of *Bmal1*^flox/flox^ mice 4 weeks before performing a SNC at ZT20 and ZT8 (**Figure 4E**). Target re-innervation was assessed 3 weeks after SNC by measuring the number of epidermal sensory fibres present in the hind paws and by measuring the number CTB positive GFP infected DRG neurons, where CTB was injected in the gastrocnemius and tibialis anterioris muscles 1 week prior to sacrificing the mice (**Figure 4E**). Remarkably, in line with our previous data, we found that 21 days post-SNC, target re-innervation in the skin (**Figure 4F-G**) and in the muscles (**Figure 4H-I**) were augmented when SNC was performed at ZT20 compared to ZT8 and that this effect was abolished in mice lacking functional Bmal1 in DRG neurons (**Figure 4F-I**).

These data indicate that Bmal1 deletion hinders the ability of DRG neurons to mount a regenerative response in a time-of-day dependent manner as shown by the reduced expression and activation of RAGs as well as reduced levels of regenerative H3K27ac when SNC is performed at ZT20.

Together, they suggest a crucial role for Bmal1 in orchestrating the transcriptional and signalling events underpinning the regenerative response to injury of DRG sensory neurons.

### Lithium treatment promotes axonal regeneration after sciatic nerve injury via a clock-dependent mechanism

We have thus far demonstrated that DRG neurons require a functional oscillating clock to be able to mount an appropriate regenerative response. Indeed, we found that regeneration is increased at specific times-of-day and that this increase is impaired by disrupting the clock *in vivo*, suggesting that manipulating the clock may promote axon regeneration. Interestingly, recent studies showed that chronoactive drugs, such as lithium, are able to interfere with the clock modulating the amplitude of circadian rhythms^33^. Lithium is widely used as a treatment of bipolar disorder, a neurological disorder associated with disrupted circadian rhythms^17–19^ and associated with increased expression of clock genes, including *Bmal1*^34–36^. Mechanistically, lithium has been suggested to act also on glycogen synthase kinase-3β (GSK-3β), which regulates degradation of CRY2, a canonical clock protein determining circadian period^17,33–36^. Additionally, *Gsk-3β* is a key regeneration-associated gene whose inhibition has been shown to increase axonal regeneration after injury^37,38^. These observations suggest the hypothesis that lithium could promote axonal regeneration. To test this, we performed SNC at ZT8 (with lower regenerative potential) in WT vs *Bmal1*^-/-^ neurons and *Cry1/2*^-/-^ mice and assessed regeneration 72h later (**Figure 5A**). We first treated mice by injecting lithium chloride i.p. (1 mEq/Kg) right after injury, then lithium carbonate (600 mg/L) was given in drinking water for 3 days from the day of surgery until sacrifice. Remarkably, we found that lithium can promote regeneration in WT (**Figure 5B-E**) but not in *Bmal1*^-/-^ (**Figure 5B-C**) nor in *Cry1/2*^-/-^ mice (**Figure 5D-E**), suggesting that the effect of lithium is due to the presence of an intact clock. No signs of activated cell death pathways by activated caspase 3 immunostaining were detected in the *Bmal1*^-/-^ and *Cry1/2*^-/-^ DRG neurons (**Supplementary Figure 17**). Additionally, we observed that both pCREB and pS6 levels are increased by lithium treatment in WT but not in *Cry1/2^-/-^* DRG neurons after injury (**Supplementary Figure 18**). Importantly, lithium induced BMAL1 expression (**Supplementary Figure 19**) but not CREB or S6 phosphorylation (**Supplementary Figure 20**) in naïve DRG neurons, supporting the idea that lithium promotes repair by acting via the molecular clock to modulate injury-dependent regenerative pathways. Together, these data show that enhancing the amplitude of the circadian rhythm with lithium promotes nerve regeneration, paving the way to the use of chrono-active compounds for nervous system repair.

**Figure 5.**
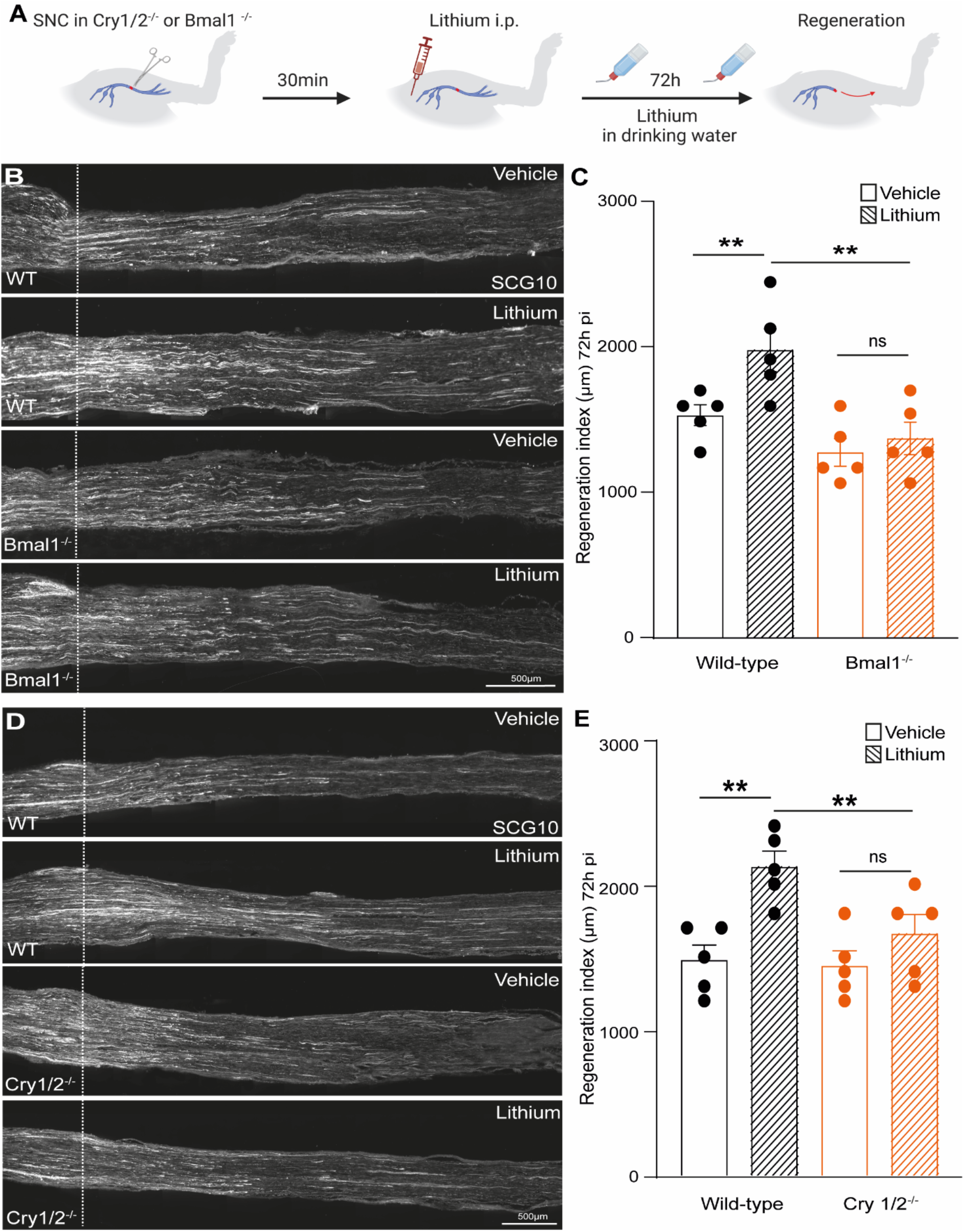
Lithium promotes axonal regeneration after SNC *in vivo.* **A.** Timeline for the *in vivo* experiment. **B.** Representative images of sciatic nerves from WT and *Bmal1*^-/-^ DRG neurons treated with lithium or vehicle and stained for SCG10 72h after SNC. Scale bar, 500 µm. **C.** Quantification of SCG10 intensity from the lesion site shown as regeneration index (distance from the injury site at which 50% of the fluorescence decays) (two-way ANOVA, Tukey’s post-hoc, p<0.05, n = 5 biologically independent animals/group). Fluorescence intensity was measured in one series of tissue sections for each nerve. **D**. Representative images of sciatic nerves from WT and *Cry1/2*^-/-^ animals treated with lithium or vehicle and stained for SCG10 72h after SNC. Scale bar, 500 µm. **E.** Quantification of SCG10 intensity from the lesion site shown as regeneration index (distance from the injury site at which 50% of the fluorescence decays) (two-way ANOVA, Tukey’s post-hoc, p<0.05, n = 5 biologically independent animals/group). Fluorescence intensity was measured in one series of tissue sections for each nerve.

## Discussion

In this study, we report evidence for a functional role of the circadian clock machinery in DRG neurons that regulates the timing and extent of axonal regeneration following a sciatic nerve injury. This circadian clock relies on the Bmal1-dependent neuronal intrinsic regulation of the regenerative ability of DRG sensory neurons. Specifically, we found that a SNC injury performed at ZT20 induces a regenerative transcriptional program dependent upon circadian rhythm, which elicits a long-lasting increase in regeneration and target re-innervation compared to an injury performed at other ZTs. Importantly, this effect was abolished when we disrupted the clock in DRG neurons by conditionally deleting the non-redundant clock core protein BMAL1. Furthermore, we observed an endogenous, oscillating clock in DRG neurons, which was disrupted by the time of the day specific deletion of *Bmal1*. Lastly, we identified lithium, a chrono-active drug widely used to treat bipolar disorder, as a proof of principle that pharmacological manipulation of the circadian clock might be a new avenue to promote repair after injury, although more systematic work will be needed to exploit its full potential.

Inflammatory cells and homeostatic processes possess circadian features and previous studies reported circadian oscillation affecting inflammatory cell numbers and activation of homeostatic conditions ^13,14^. However, here we did not observe post-injury diurnal changes in macrophages, neutrophils or Schwann cells infiltration at the sciatic injury site. This is in line with our findings supporting a neuronal intrinsic molecular clock regenerative mechanism priming DRG neurons in a time-of-day dependent manner. In fact, our data indicate a clock-dependent priming of DRG neurons prior to the injury where there is no inflammation or leukocyte infiltration, suggesting that the role of non-neuronal cells in clock-dependent reprogramming of DRG neurons may be of secondary importance in the circadian control of nerve regeneration.

Evidence of an interplay between an injury response and circadian system has been reported in a number of tissues and models including skin wound healing^16^, intestinal regeneration^39^, spinal cord injury ^40,41^ and pain^21^. However, the specific circadian regulation of the axonal regenerative ability has remained elusive so far. Here we show that Bmal1 is required for sciatic nerve regeneration and target re-innervation as well as for the activation of the regenerative program of DRG sensory neurons. Bmal1 neuronal conditional deletion was able to selectively impair the regenerative response of DRG neurons at the regeneration peak (ZT20) while no significant differences in regeneration or activation of RAGs and RAEs were found at the regeneration trough (ZT8). Thus, *Bmal1* acts as time-of-day dependent master regulator of RAGs and RAEs by regulating transcription factors and histone acetylation, ultimately coordinating and synchronising molecular mechanisms for successful axonal regeneration. The evidence that peripheral neuronal *Bmal1* deletion is sufficient to abolish time-of-day-dependent nerve regeneration suggests that the central clock is not required for this regenerative phenotype. Additionally, the *Bmal1*-dependent circadian control of regeneration does not seem to depend upon locomotor activity because neuronal *Bmal1* deletion does not disrupt locomotion.

Since manipulation of *Bmal1* expression impairs the physiological oscillation of the circadian clock, *Bmal1* overexpression may not be a feasible approach to enhance regeneration. However, the future identification of the molecular mechanisms controlling the neuronal molecular clock holds the promise to positively affect the regenerative ability after nervous system injuries.

Importantly, a recent study reported that burn injury healing time in humans is increased ∼60% when burns occurred during the resting phase (night) compared with during the active phase (day) ^16^, suggesting that the power of clock-dependent mechanisms may be harnessed via chronotherapeutic approaches, including timed therapies and chrono-active drugs ^42^. In line with this, we successfully used lithium, a chrono-active drug, as a proof of principle approach in a model of sciatic nerve injury. Finally, the circadian influence on axonal regeneration suggests that repair strategies relying on neurorehabilitation or neuronal activity should consider time-tuned approaches during the 24 hours cycle to align with changes in the neuronal intrinsic regenerative ability.

Given the pervasive nature of circadian clocks, our findings finally suggest a careful re-evaluation of the data collected thus far in the field that could have been affected by the different time-of-day at which the experiments have been performed. It also calls for caution in designing future experiments, which must hereafter consider clock-dependent mechanisms.

### Limitations of the study

The study provides a first evidence of a functional clock in DRG sensory neurons that controls time-of-day dependent regenerative ability by regulating the transcriptional landscape of DRG neurons. Additionally, it provides initial evidence of the possibility to exploit clock-dependent mechanisms to implement or develop future clinically suitable strategies. However, it also bears some limitations. i) while we demonstrate that Bmal1 is required for the gene expression of RAGs supporting nerve regeneration, due to intrinsic technical limitations of the model (low DNA yield from DRG for successful Cut&Run), we were not able to provide direct evidence that BMAL1 occupies gene regulatory elements of these genes; ii) due to current unavailability of specific chrono-active compounds, we used lithium as chrono-active drug, which, while of high translational value, it has pleiotropic effects; (iii) this study was performed in a model of peripheral nerve injury, and we have no evidence of whether enhancing the amplitude of the molecular clock could benefit axon regeneration in a model of central nervous system injury.

## Acknowledgments

This work was supported by start-up funds from the Department of Brain Sciences, Imperial College London (SDG); The Rosetrees Trust (SDG). The research was supported by the National Institute for Health Research (NIHR), Imperial Biomedical Research Centre (MED, SDG). The research was supported by the National Institute for Health Research (NIHR) Imperial Biomedical Research Centre (SDG). The views expressed are those of the author(s) and not necessarily those of the NHS, the NIHR or the Department of Health.

## Author contributions

F.D.V. designed, performed experiments, data analysis and wrote the manuscript; F.M. performed data analysis; I.P. performed experiments and data analysis, J.S.C. performed experiments; L.L.G. performed experiments and data analysis; A.G. performed experiments and data analysis; Y.Y. performed experiments; M.C.D. performed data analysis; C.P.M. performed experiments; L.Z. performed experiments; G.K. performed experiments; E.S. performed experiments; T.H.H. performed experiments; C.S. designed experiments and edited the manuscript M.B. designed experiments and edited the manuscript; S.D.G. designed experiments, provided funding and edited the manuscript.

## Data availability

The datasets analysed during the current study are available in the following repository: GSE161342; GSE138769; GSE97090; GSE224316; GSE235687.

---

**Supplementary Figure 1.**
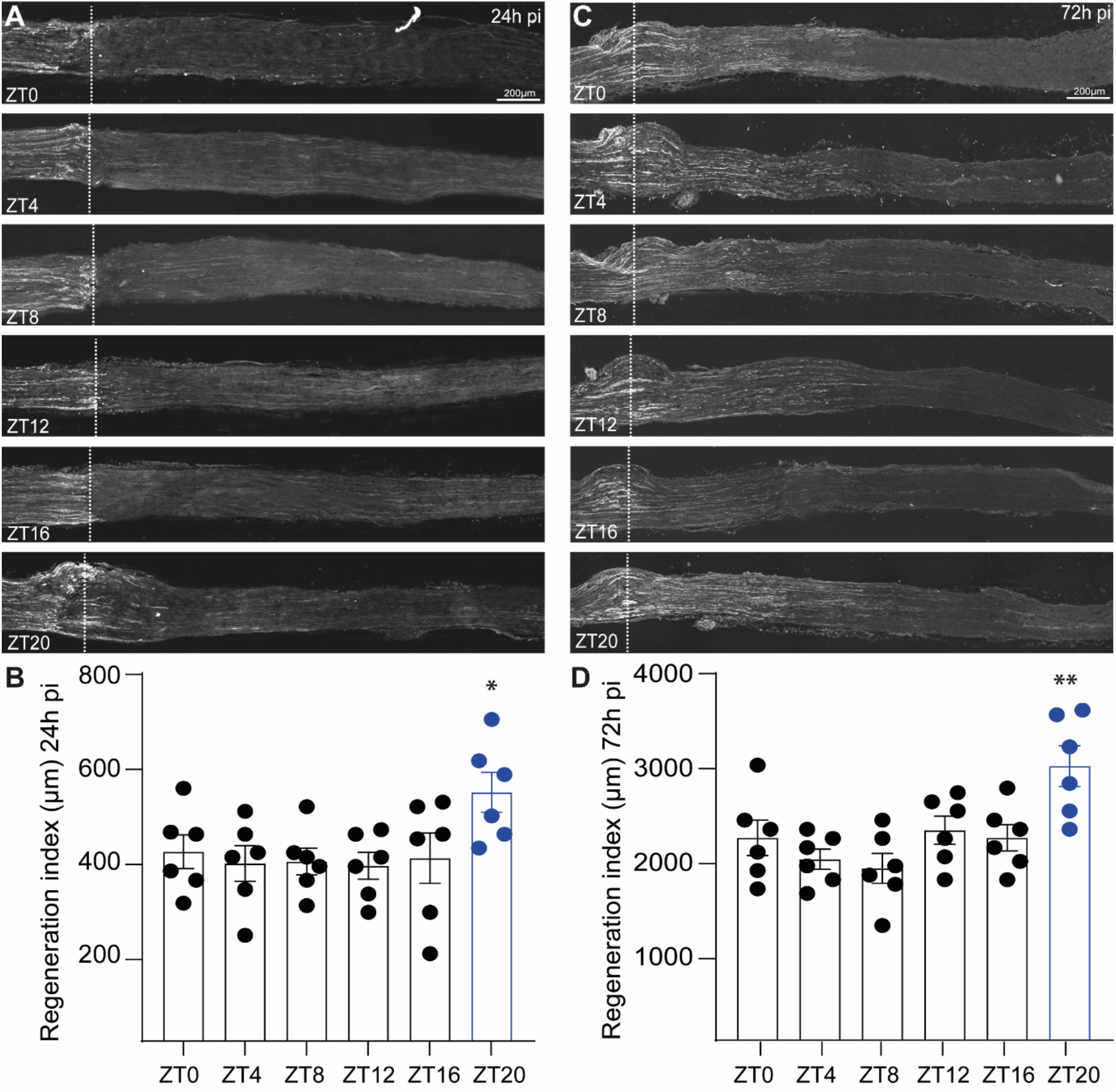
**A.** Representative images of sciatic nerves injured at ZT8 or ZT20 stained for SCG10 24h after SNC. Scale bar, 500 µm. **B.** Quantification of SCG intensity from the lesion site shown as regeneration index 24h after injury (distance from the injury site (dotted lines) at which 50% of the fluorescence decays) (mean ± SEM, one-way ANOVA, Tukey’s post-hoc, p<0.05, n = 6 biologically independent animals/group). Fluorescence intensity was measured in one series of tissue sections for each nerve. **C.** Representative images of sciatic nerves injured at ZT8 or ZT20 stained for SCG10 72h after SNC. Scale bar, 500 µm. **D.** Quantification of SCG intensity from the lesion site shown as regeneration index 72hr after injury (distance from the injury site (dotted lines) at which 50% of the fluorescence decays) (mean ± SEM, one-way ANOVA, Tukey’s post-hoc, p<0.05, n = 6 biologically independent animals/group). Fluorescence intensity was measured in one series of tissue sections for each nerve.

**Supplementary Figure 2.**
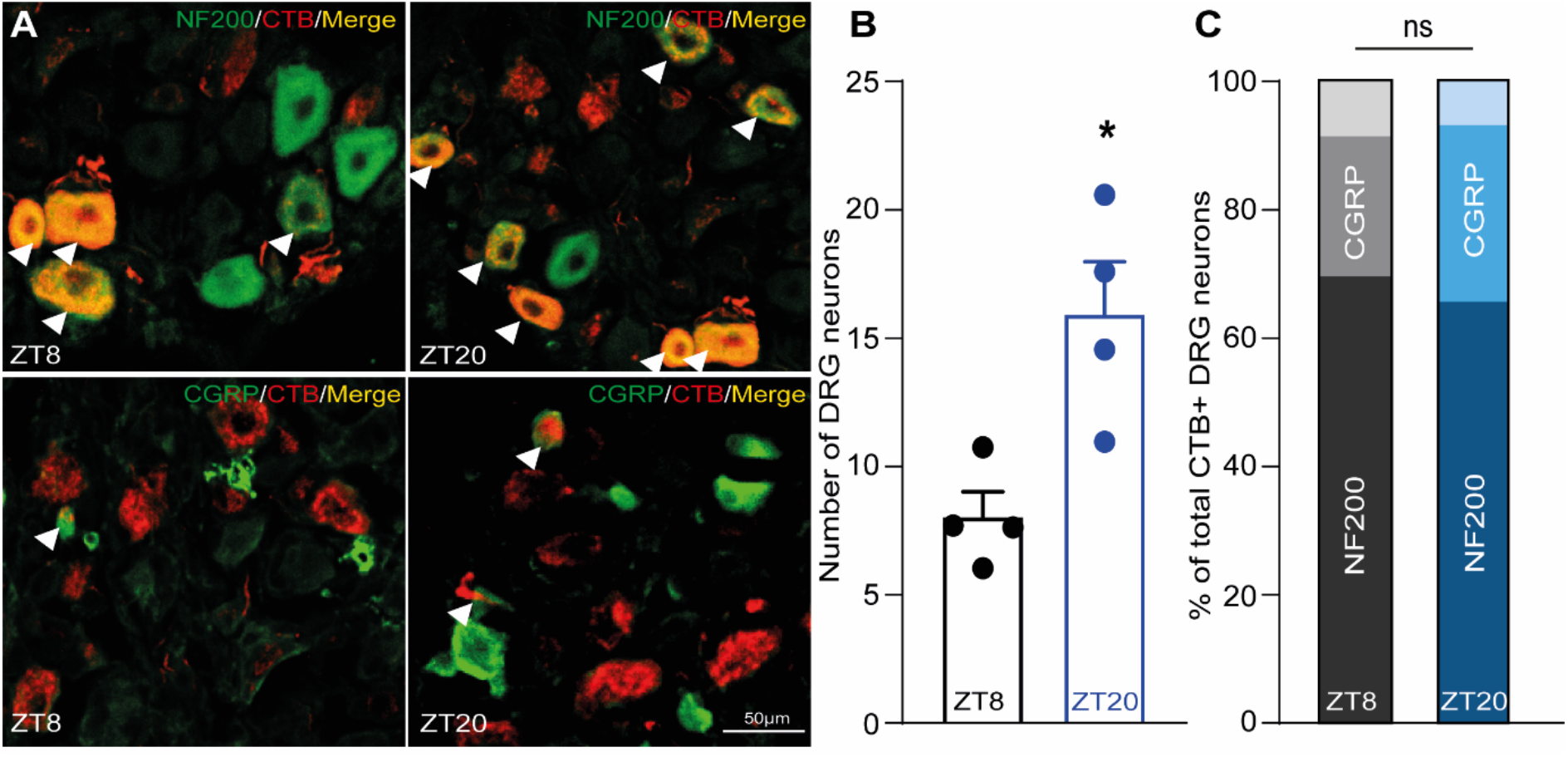
**A.** Representative images of DRG injured at ZT8 or ZT20, injected with CTB retrograde tracer 429 (red), stained with NF-200 (green, top) or CGRP (green, bottom) and analysed 3 days afterSNC. Arrowheads highlight double positive DRG neurons (yellow). Scale bar, 50 µm. **B.** Quantification of CTB positive DRG neurons injured at ZT8 or ZT20 (Student T-test, p<0.05, n = 4 biologically independent animals/group). Number of cells was measured in one series of tissue sections for each DRG. **C.** Quantification of CTB and NF-200 (or CGRP) double positive DRG neurons injured at ZT8 or ZT20 (Student T-test, p<0.05, n = 4 biologically independent animals/group). Number of cells was measured in one series of tissue sections for each DRG.

**Supplementary Figure 3.**
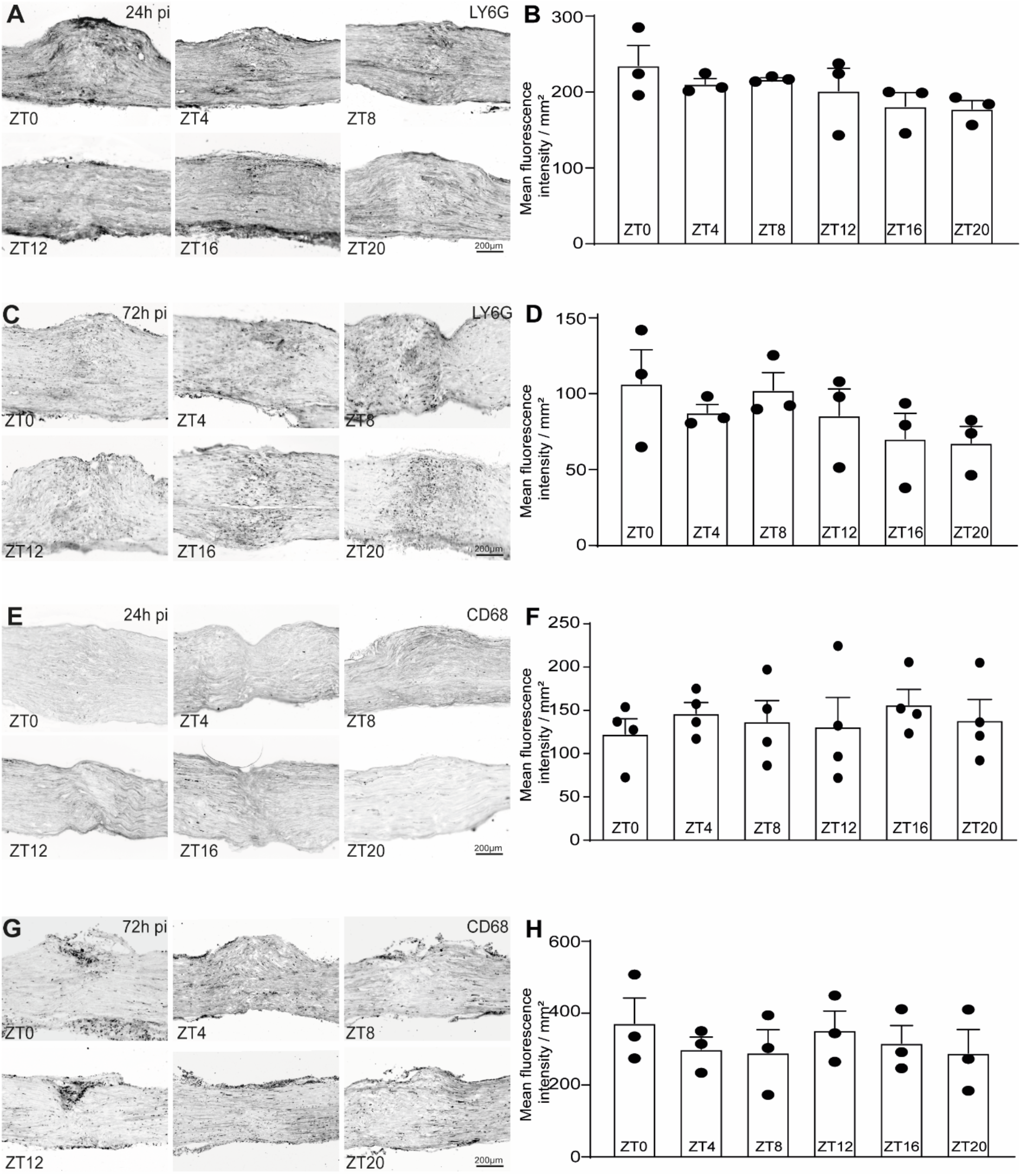
**A.** Representative images of sciatic nerves from animals injured at ZT0, ZT4, ZT8, ZT12, ZT16 or ZT20 stained for LY6G 24h after SNC. Scale bar, 200 µm. **B.** Quantification of LY6G fluorescence intensity from the lesion site 24h after SNC (mean ± SEM, one-way ANOVA, Tukey’s post-hoc, p<0.05, n = 4 biologically independent animals/group). Fluorescence intensity was measured in one series of tissue sections for each nerve. **C.** Representative images of sciatic nerves from animals injured at ZT0, ZT4, ZT8, ZT12, ZT16 or ZT20 stained for LY6G 72h after SNC. Scale bar, 200 µm **D.** Quantification of Lys6G fluorescence intensity from the lesion site 72h after SNC (mean ± SEM, one-way ANOVA, Tukey’s post-hoc, p<0.05, n = 4 biologically independent animals/group). Fluorescence intensity was measured in one series of tissue sections for each nerve. **E.** Representative images of sciatic nerves from animals injured at ZT0, ZT4, ZT8, ZT12, ZT16 or ZT20 stained for CD68 24h after SNC. Scale bar, 200 µm. **F.** Quantification of CD68 fluorescence intensity from the lesion site 24h after SNC (mean ± SEM, one-way ANOVA, Tukey’s post-hoc, p<0.05, n = 4 biologically independent animals/group). Fluorescence intensity was measured in one series of tissue sections for each nerve. **G.** Representative images of sciatic nerves from animals injured at ZT0, ZT4, ZT8, ZT12, ZT16 or ZT20 stained for CD68 72h after SNC. Scale bar, 200 µm **H.** Quantification of CD68 fluorescence intensity from the lesion site 72h after SNC (mean ± SEM, one-way ANOVA, Tukey’s post-hoc, n = 4 biologically independent animals/group). Fluorescence intensity was measured in one series of tissue sections for each nerve.

**Supplementary Figure 4.**
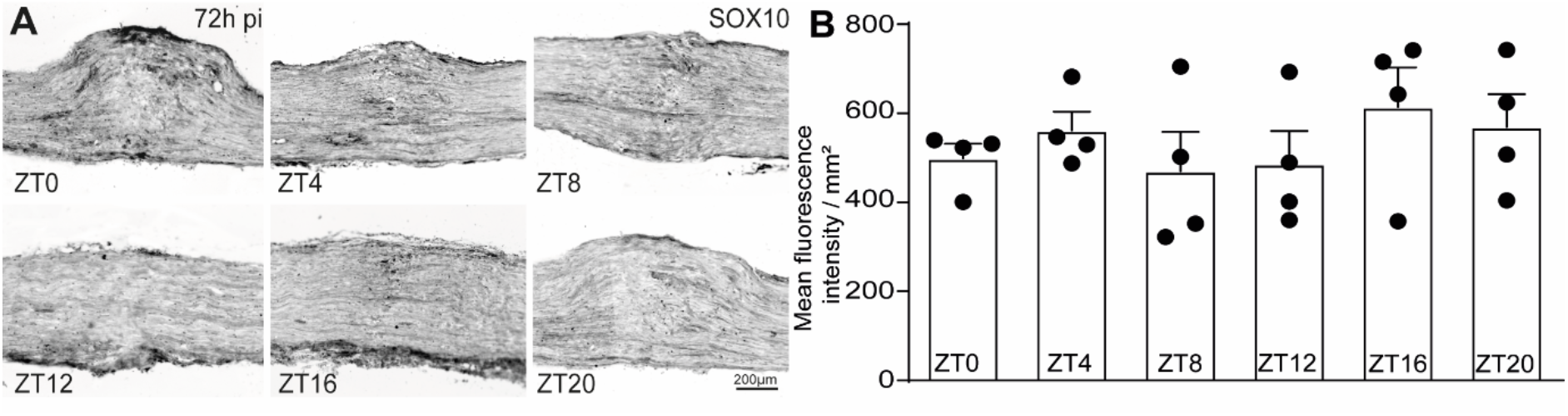
**A.** Representative images of sciatic nerves from animals injured at ZT0, ZT4, ZT8, ZT12, ZT16 or ZT20 stained for SOX10 72h after SNC. Scale bar, 200 µm. **B.** Quantification of SOX10 fluorescence intensity from the lesion site 72h after SNC (mean ± SEM, one-way ANOVA, Tukey’s post-hoc, p<0.05, n = 4 biologically independent animals/group). Fluorescence intensity was measured in one series of tissue sections for each nerve.

**Supplementary Figure 5.**
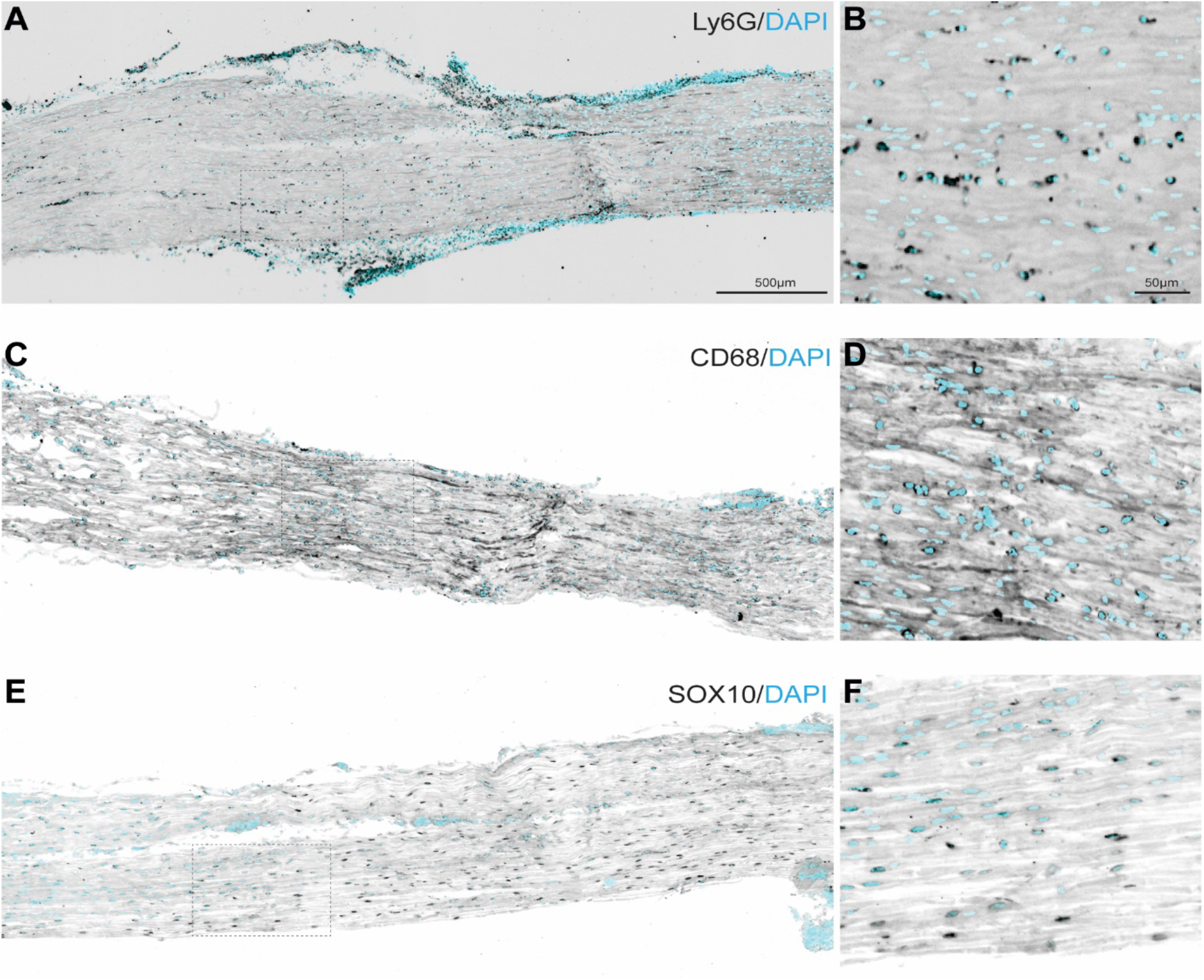
**A.** Representative pictures of the sciatic nerve after injury immunostained for Neutrophils (Ly6G, black) and DAPI (cyan). Scale bar 500μm. B. Close-up magnification of the dotted highlighted area in A. Scale bar 50μm **A**. **C.** Representative pictures of the sciatic nerve after injury immunostained for Macrophages (CD68, black) and DAPI (cyan). Scale bar 500μm **D.** Close-up magnification of the dotted highlighted area in C. Scale bar 50μm**. E.** Representative pictures of the sciatic nerve after injury immunostained for Schwann cells (SOX10, black) and DAPI (cyan). Scale bar 500μm **F.** Close-up magnification of the dotted highlighted area in E. Scale bar 50μm.

**Supplementary Figure 6.**
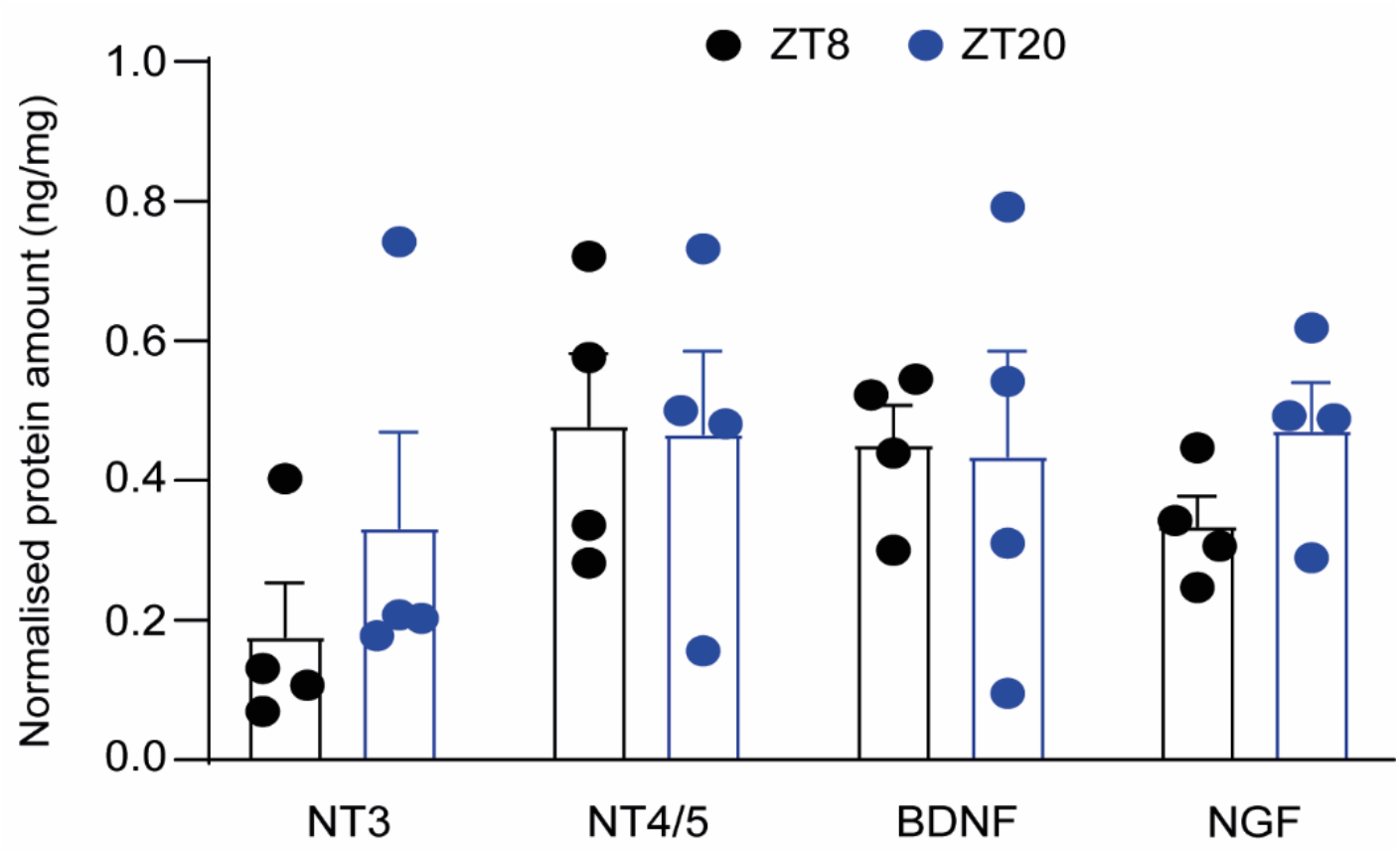
Quantification of neurotrophins levels in whole DRG lysate (mean ± SEM, Two-way ANOVA, Sidak’s post-hoc, p<0.05, n = 4 biologically independent animals/group).

**Supplementary Figure 7.**
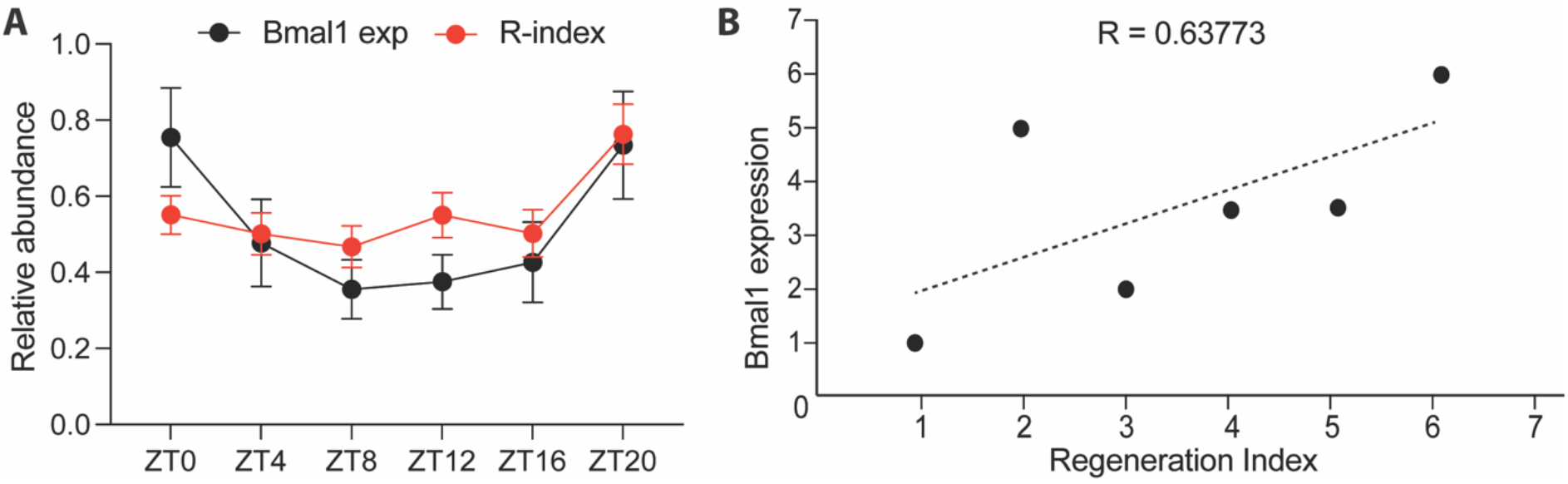
**A.** Overlapped data points of quantitative RT-PCR analysis of the mRNA levels of Bmal1 mRNA levels in DRG at ZT8, ZT12, ZT16, ZT20, ZT24 ZT0 and ZT4, and normalised over GAPDH (Figure 1C, black line) regeneration index (R-index) (Figure 1D **and Supplementary** Figure 1, red line). **B.** Scatter plot representing positive correlation between Bmal1 mRNA expression (Figure 1C) vs Regeneration Index (Figure 1D **and Supplementary** Figure 1) calculated as Pearson correlation coefficient (R) (N=5).

**Supplementary Figure 8.**
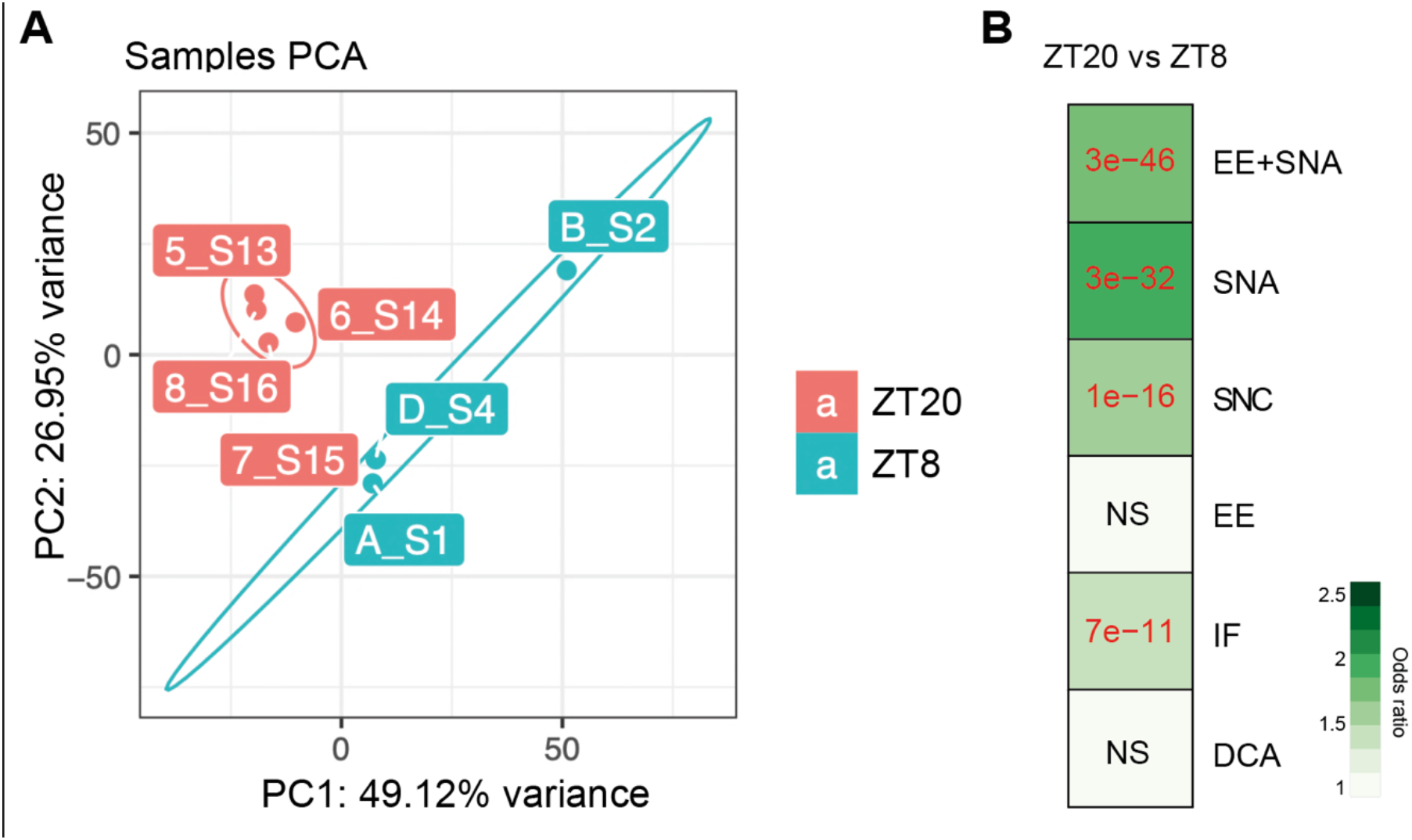
**A**. PCA analysis of all experimental groups (ZT20 vs ZT8) clustered according to percentage of variance (A, B, D=ZT8; 5, 6, 7, 8=ZT20). B. Odds ratio analsyis of the RNA-seq dataset against previously published datasets in DRG (as referred to in Figure 1B) (in red adjusted p-value<0.05, NS=not significant; EE=Enriched environment, SNA=sciatic nerve axotomy, SNC=Sciatic nerve crush, IF=intermittent fasting, DCA=dorsal column axotomy).

**Supplementary Figure 9.**
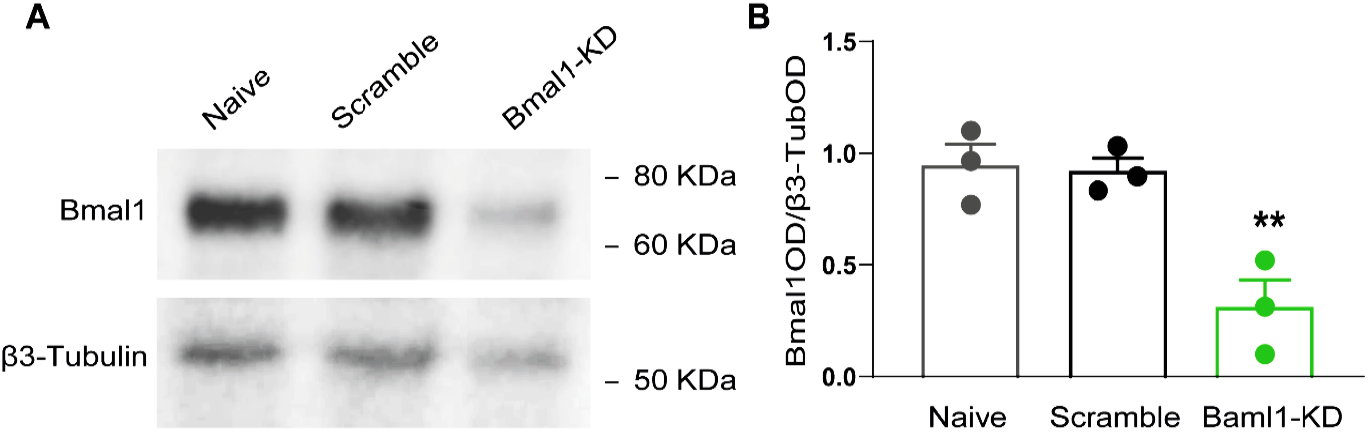
**A**. Immunoblotting analysis of BMAL1 from protein extracts of neuronal enriched DRG cultures. **B.** Quantification of immunoblotting, βIII-tubulin was used as a loading control to which protein expression was normalized. (mean ± SEM, one-way ANOVA, Tukey’s post hoc, p<0.05, n = 3 biologically independent animals/group examined over three independent experiments).

**Supplementary Figure 10.**
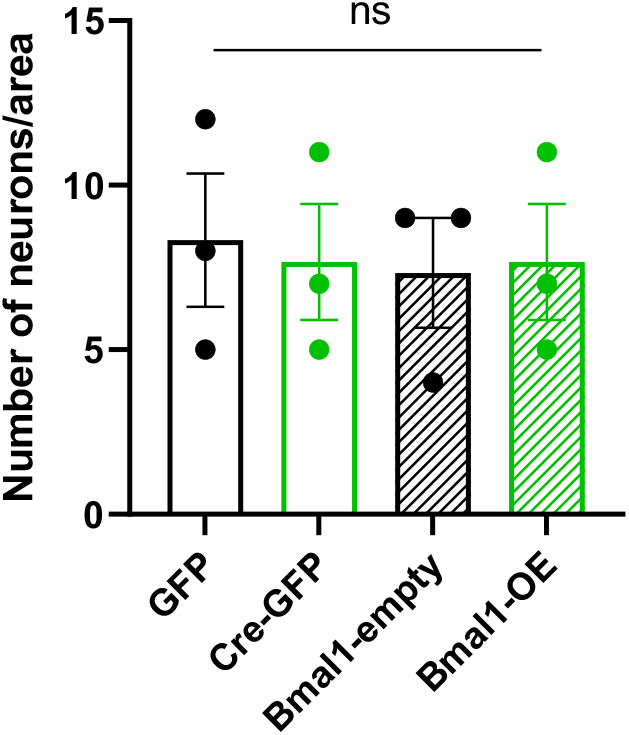
Quantification of neurons/area in cultures quantified by DAPI in control AAV-GFP, AAV-Cre-GFP, AAV-Cre-GFP+Bmal1-empty vector and AAV-Cre-GFP+Bmal1-OE vector groups (as in Figure 3D-G). (One-way ANOVA Tukey’s post-hoc, ns=not significant, n = 3).

**Supplementary Figure 11.**
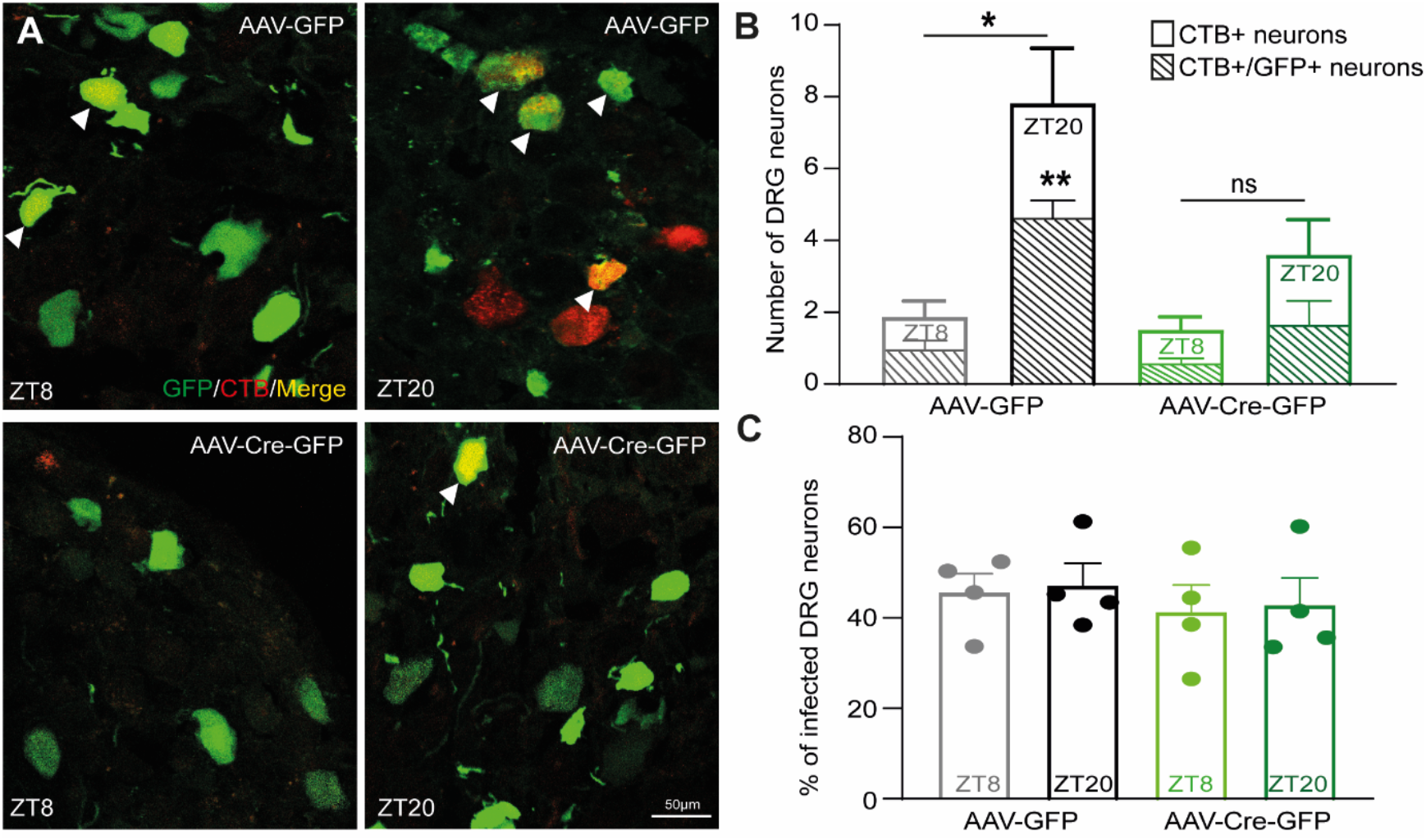
**A.** Representative images of DRGs from Bmal1^fl/fl^ animals infected with GFP vs Cre-GFP virus (green) injured at ZT8 or ZT20, injected with CTB retrograde tracer (red) and analysed 3 days after SNC. Arrowheads highlight double positive DRG neurons (yellow). Scale bar, 50 µm. **B.** Quantification of CTB and GFP double positive DRG neurons (mean ± SEM, two-way ANOVA, Tukey’s post hoc, p<0.05, n = 4 biologically independent animals/group). Number of cells was measured in one series of tissue sections for each DRG. **C.** Quantification of transduction across samples shown as percentage of infected DRG neurons (mean ± SEM, two-way ANOVA, Tukey’s post hoc, n = 4 biologically independent animals/group). Number of cells was measured in one series of tissue sections for each DRG.

**Supplementary Figure 12.**
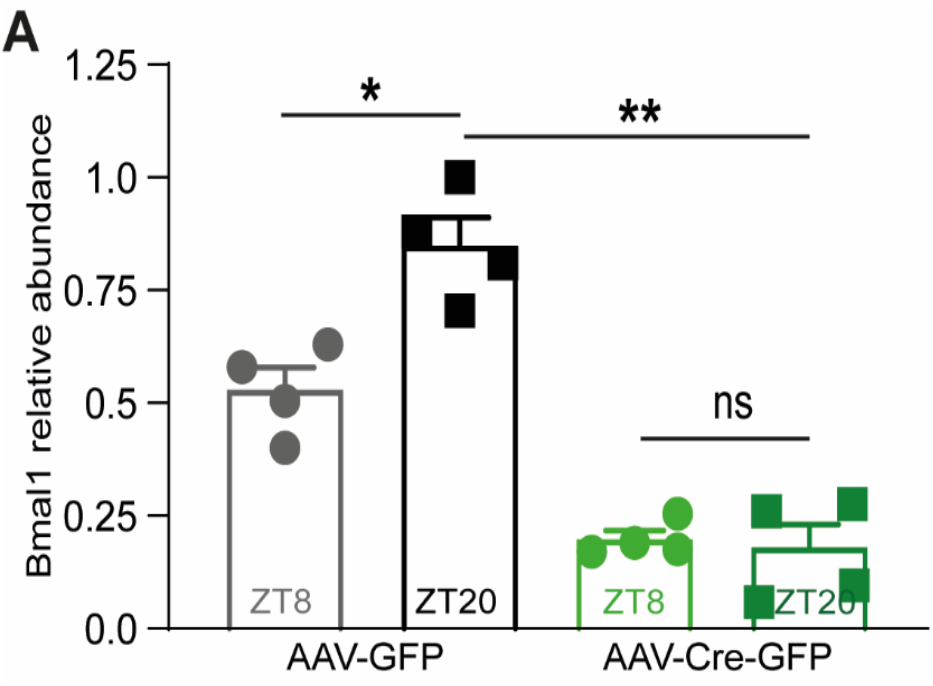
Quantitative RT-PCR analysis of the mRNA levels of *Bmal1* normalised over *Gapdh* (mean ± SEM, two-way ANOVA, Tukey’s post hoc, p<0.05, n=4 biologically independent animals/group examined over 3 independent experiments).

**Supplementary Figure 13.**
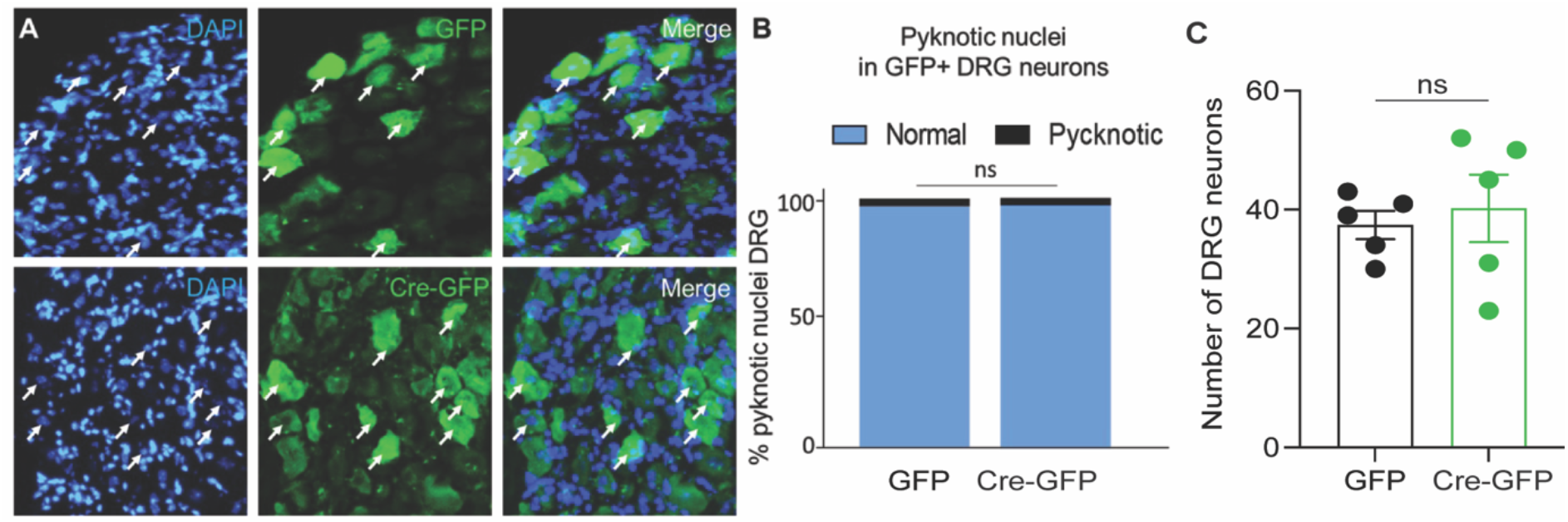
**A.** Representative images of DRGs from Bmal1^flox/flox^ animals infected with GFP vs Cre-GFP (green) virus injured at ZT8 or ZT20. Nuclei are stained with DAPI and analysed 3 days after SNC. **B.** Quantification of pyknotic nuclei by DAPI staining in DRG (Student t-test, ns=not significant, n = 3). Number of nuclei was measured in one series of tissue sections for each DRG. **C.** Quantification of pyknotic nuclei by DAPI staining in DRG (Student t-test, ns=not significant, n = 3). Number of neurons was measured in one series of tissue sections for each DRG.

**Supplementary Figure 14.**
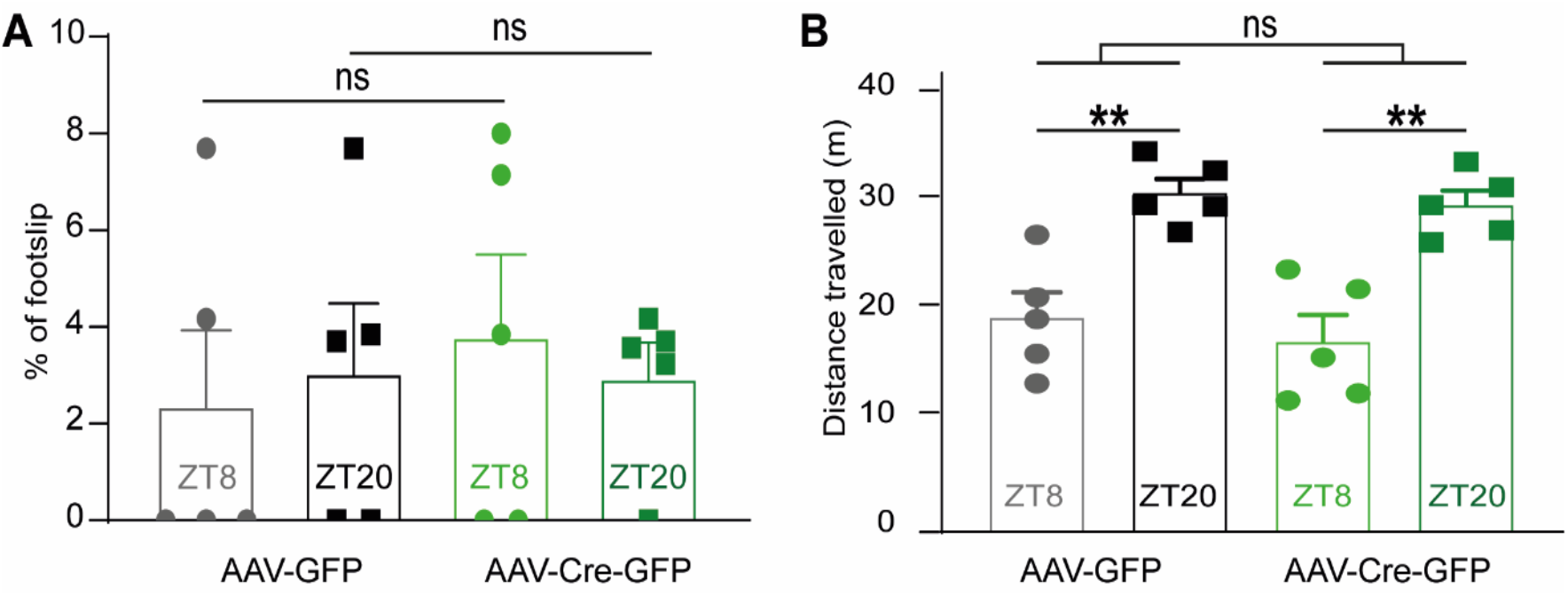
**A.** Quantification of the percentage of footslips per run on a grid walk (mean ± SEM, two-way ANOVA, Tukey’s post hoc, p<0.05, n = 5 biologically independent animals/group examined over two independent experiments). **B.** Quantification of the distance travelled (in m) in a 5min open field test (mean ± SEM, two-way ANOVA, Tukey’s post hoc, p<0.05, n = 5 biologically independent animals/group examined over two independent experiments).

**Supplementary Figure 15.**
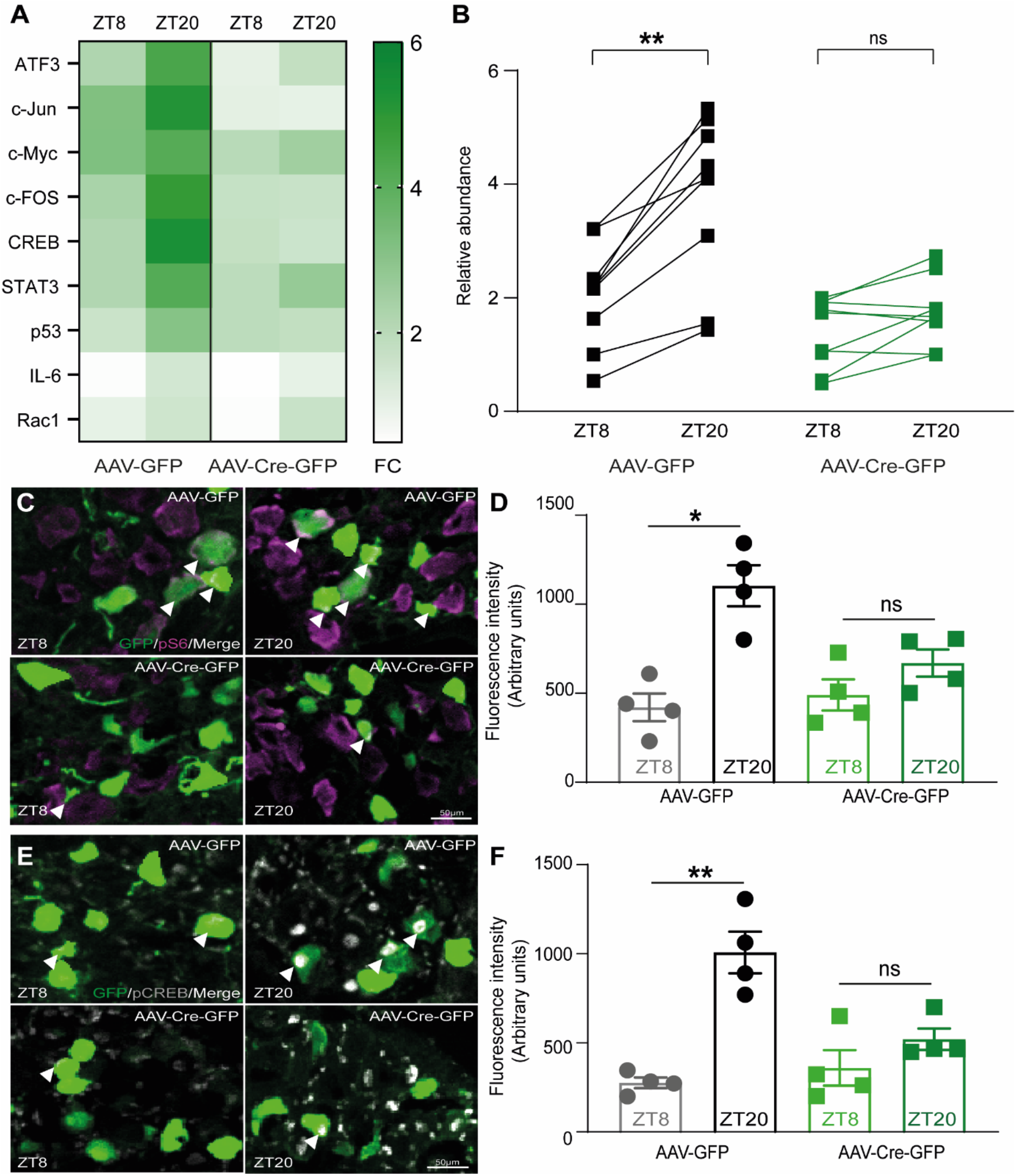
**A.** Heatmap representation of quantitative RT-PCR analysis of the mRNA levels of *ATF3*, *c-Jun*. *c-Myc*, *c-Fos*, *CREB*, *STAT3*, *p53*, *IL-6*, *Rac1* normalised over GAPDH in DRG after injury at ZT8 and ZT20 in GFP and Cre-GFP injected Bmal1^fl/fl^ mice (FC = Fold Change, n=3 biologically independent animals/group examined over 3 independent experiments). **B.** Representation of time-of-day expression levels difference of quantitative RT-PCR analysis in DRG after injury between GFP and Cre-GFP injected Bmal1^fl/fl^ mice (mean per gene, lines highlight the dynamics of the expression levels of each gene from ZT8 to ZT10 in GFP and Cre-GFP groups (mean, two-way ANOVA, Tukey’s post-hoc, p<0.05, n=3 biologically independent animals/group examined over 3 independent experiments). **C**. Representative images of DRG immunostained for pS6 (purple) and GFP (green) at ZT8 and ZT20 in GFP and Cre-GFP injected Bmal1^fl/fl^ mice. Arrowheads highlight double positive DRG neurons (white). Scale bar, 50 µm. **D.** Quantification of the fluorescence intensity of pS6 (purple) in GFP positive DRG neurons (mean ± SEM, two-way ANOVA, Tukey’s post-hoc, p<0.05, n=4 biologically independent animals/group). Fluorescence intensity was measured in one series of tissue for each DRG). **E**. Representative images of DRG immunostained for pCREB (greys) and GFP (green) at ZT8 and ZT20 in GFP and Cre-GFP injected Bmal1^fl/fl^ mice. Arrowheads highlight double positive DRG neurons (white). Scale bar, 50 µm. **F.** Quantification of the fluorescence intensity of pCREB (grey) in GFP positive DRG neurons (mean ± SEM, two-way ANOVA, Tukey’s post-hoc, p<0.05, n=4 biologically independent animals/group). Fluorescence intensity was measured in one series of tissue for each DRG.

**Supplementary Figure 16.**
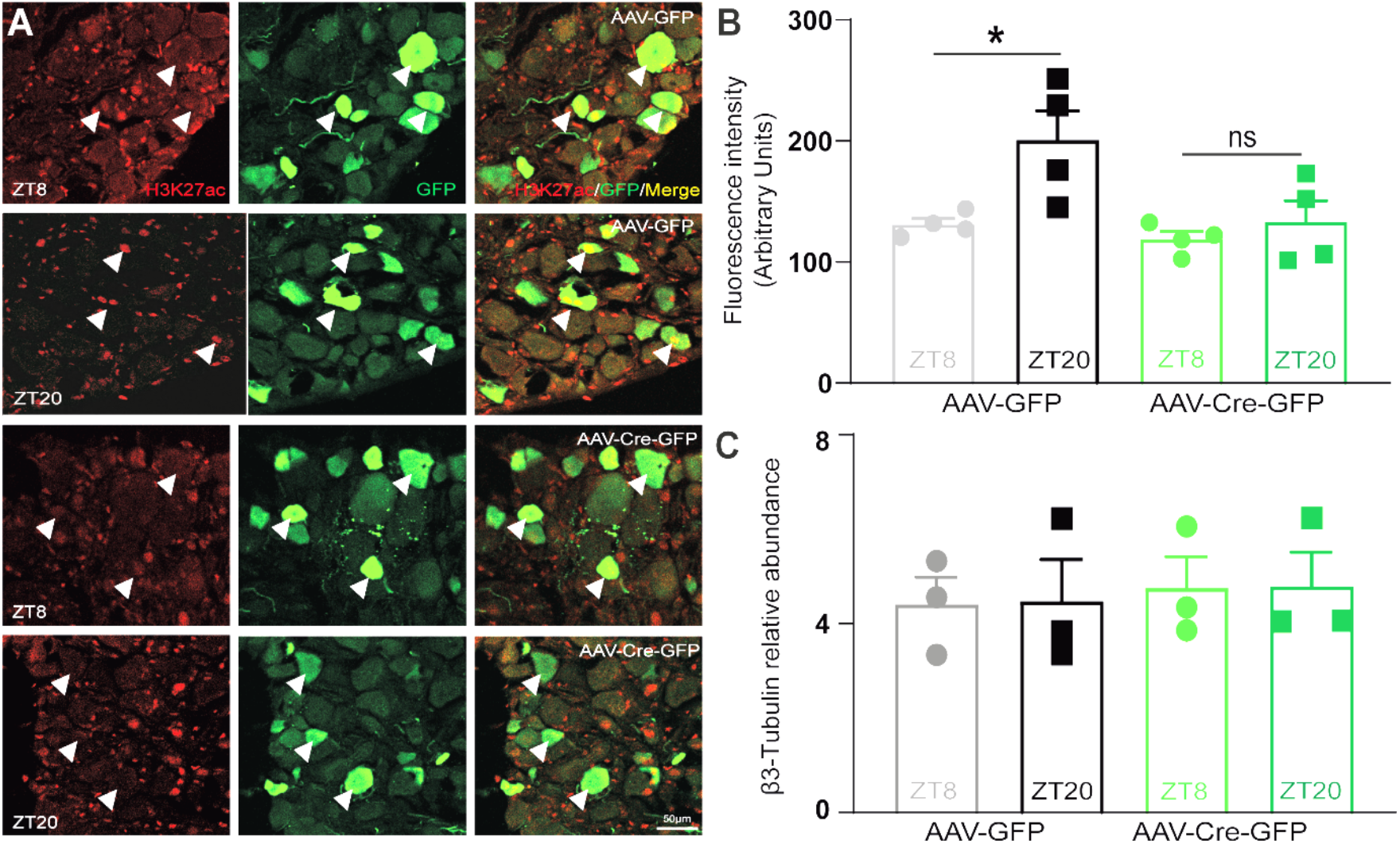
**A**. Representative images of DRG immunostained for H3K27ac (red) and GFP (green) at ZT8 and ZT20 in GFP and Cre-GFP injected Bmal1^flox/flox^ mice. Arrowheads highlight double positive DRG neurons (yellow). Scale bar, 50 µm. Scale bar, 50 µm. **B.** Quantification of the fluorescence intensity of H3K27ac (red) in GFP positive (green) DRG neurons (mean ± SEM, two-way ANOVA, Tukey’s post-hoc, p<0.05, n=4 biologically independent animals/group. Fluorescence intensity was measured in one series of tissue for each DRG). **C.** Quantitative RT-PCR analysis of the mRNA levels of *βIII-tubulin* normalised over *Gapdh* (mean ± SEM, two-way ANOVA, Tukey’s post hoc, p<0.05, n=3 biologically independent animals/group examined over 3 independent experiments).

**Supplementary Figure 17.**
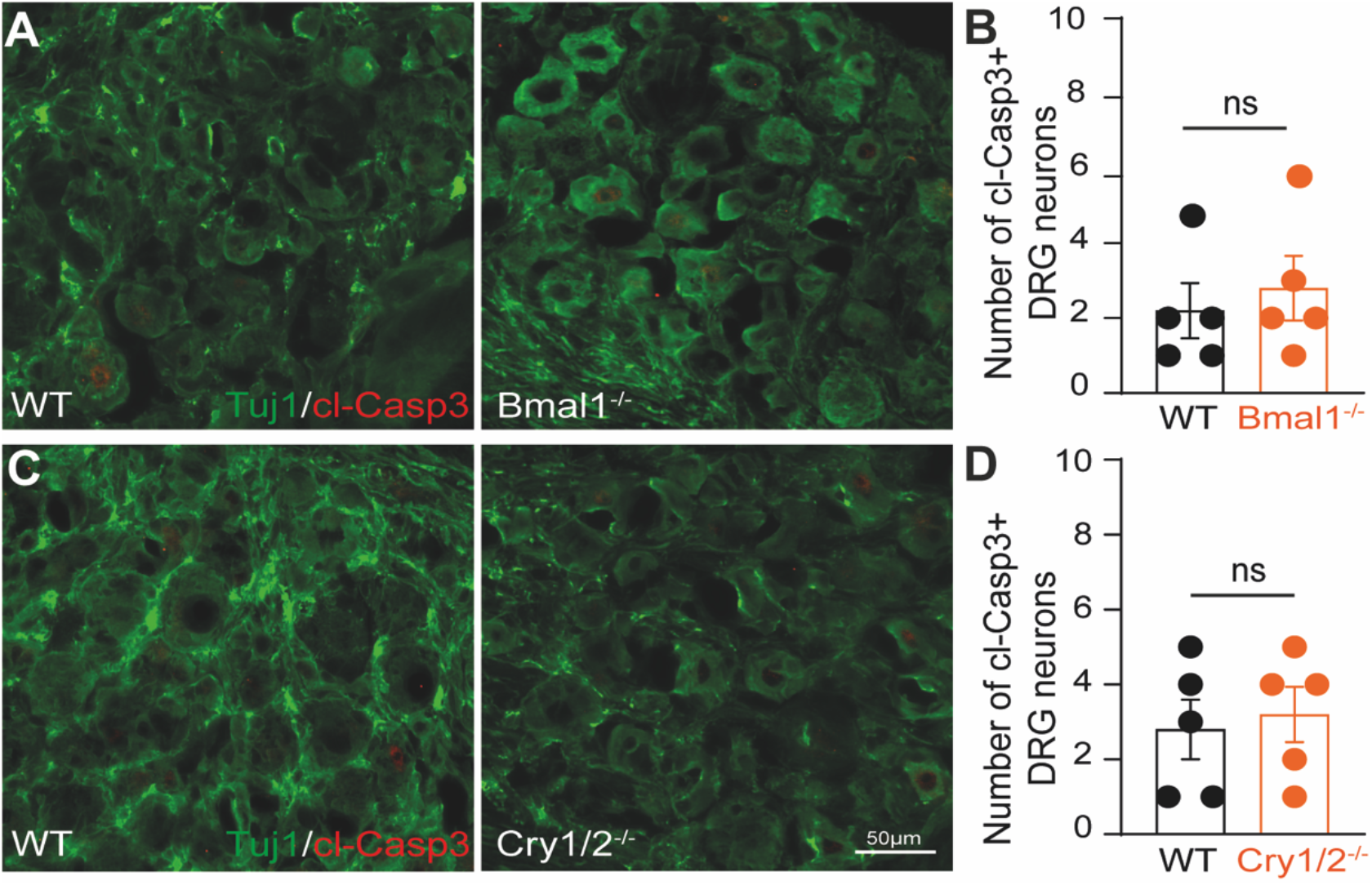
**A.** Representative images of DRGs from Bmal1**^-/-^** animals immunostained for cleaved-Caspase-3 (red) and Tuj1 (green). Scale bar, 50 µm. **B.** Quantification of cleaved-Caspase-3 positive in WT vs Bmal1**^-/-^** DRG neurons (Student T-test, ns=not significant, n = 5 biologically independent animals/group). **C.** Representative images of DRGs from Cry1/2**^-/-^** animals immunostained for cleaved-Caspase-3 (red) and Tuj1 (green). Scale bar, 50 µm. **D.** Quantification of cleaved-Caspase-3 positive WT vs Cry1/2**^-/-^** DRG neurons (Student T-test, ns=not significant, n = 5 biologically independent animals/group).

**Supplementary Figure 18.**
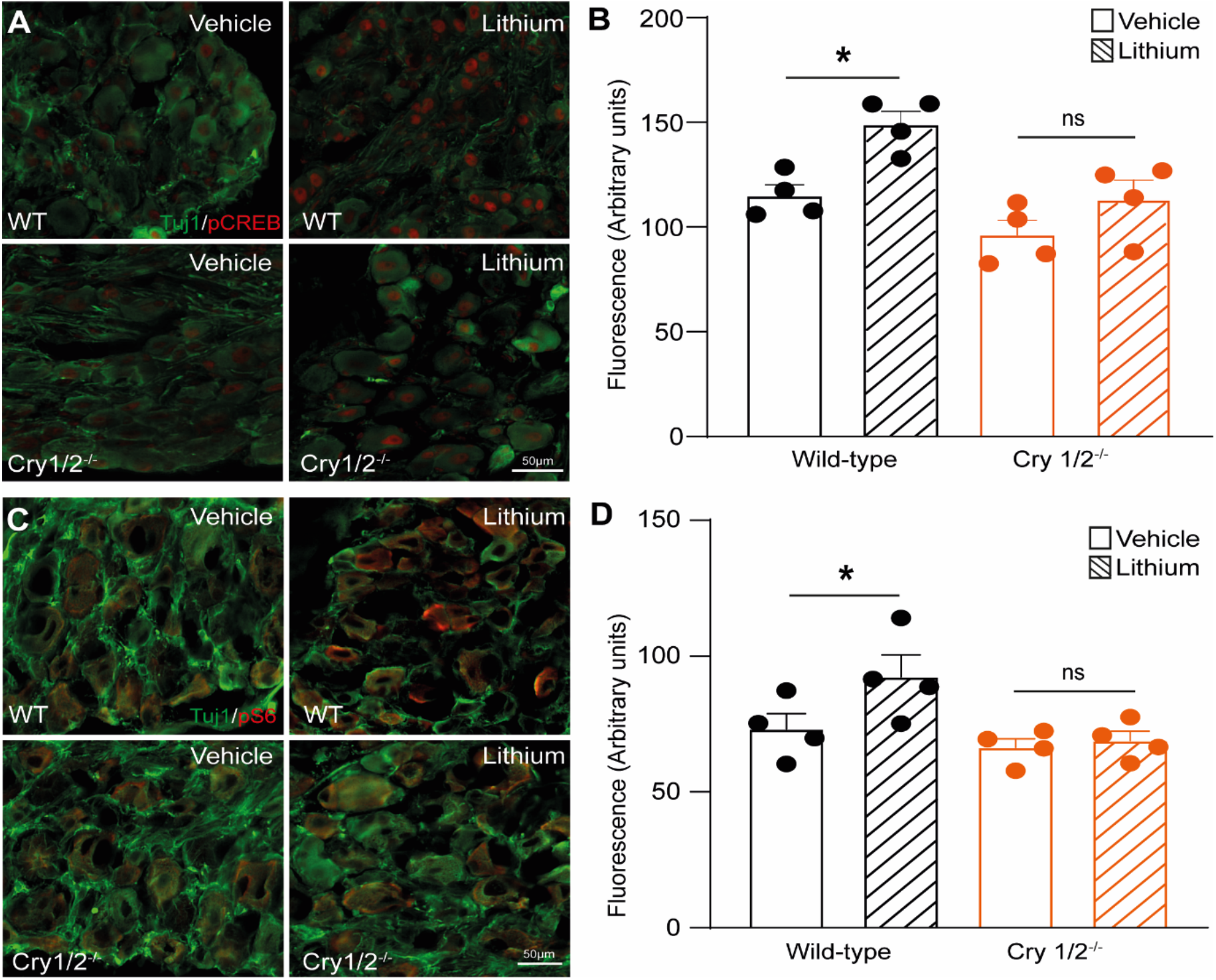
**A.** Representative images of DRG from WT and *Cry1/2*^-/-^ mice treated with lithium or vehicle after SNC immunostained for pCREB (red) and Tuj1 (green). Scale bar, 50 µm. **B.** Quantification of the fluorescence intensity of pCREB (red) in Tuj1 positive DRG neurons (mean ± SEM, two-way ANOVA, Tukey’s post-hoc, p<0.05, n=4 biologically independent animals/group). Fluorescence intensity was measured in one series of tissue for each DRG). **C**. Representative images of DRG from WT and *Cry1/2*^-/-^ mice treated with lithium or vehicle after SNC immunostained for pS6 (red) and Tuj1 (green). Scale bar, 50 µm. **D.** Quantification of the fluorescence intensity of pS6 (grey) in Tuj1 positive DRG neurons (mean ± SEM, two-way ANOVA, Tukey’s post-hoc, p<0.05, n=4 biologically independent animals/group). Fluorescence intensity was measured in one series of tissue for each DRG.

**Supplementary Figure 19.**
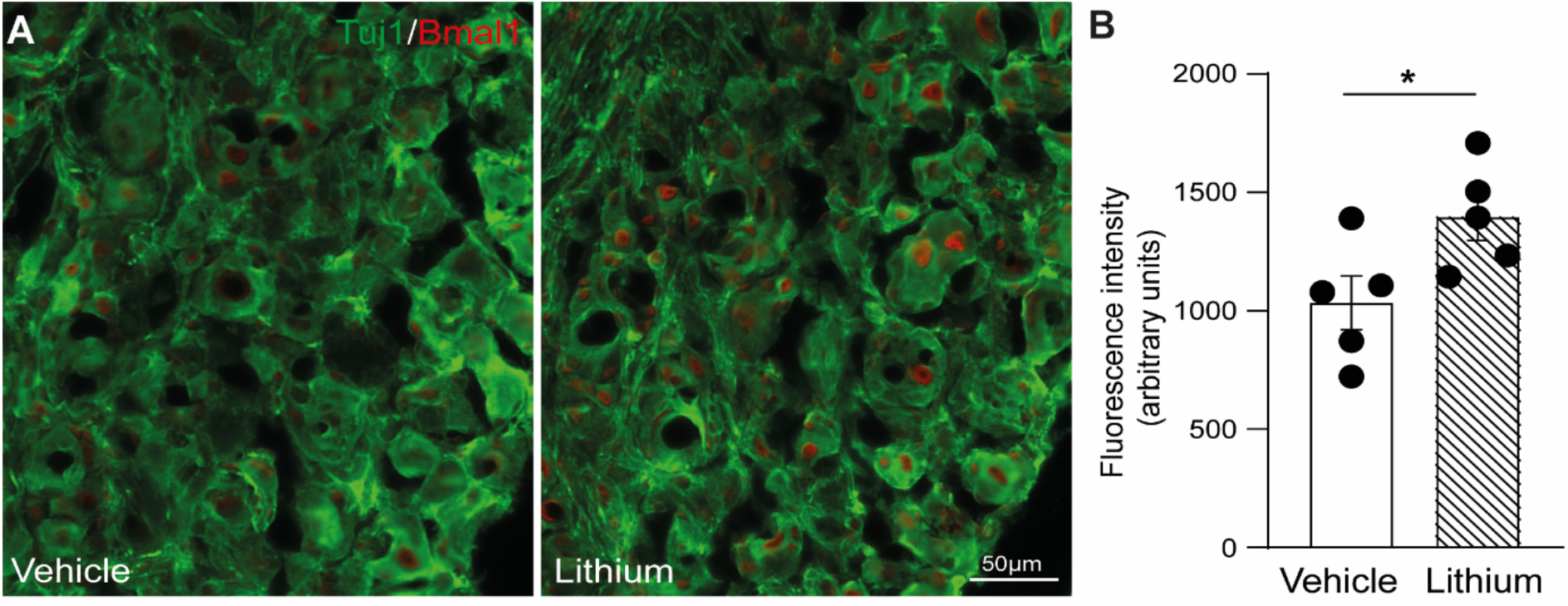
**A.** Representative images of DRG from WT mice treated with lithium or vehicle immunostained for BMAL1 (red) and Tuj1 (green). Scale bar, 50 µm. **B.** Quantification of the fluorescence intensity of BMAL1 (red) in Tuj1 positive DRG neurons (mean ± SEM, Student T-test, p<0.05, n=5 biologically independent animals/group). Fluorescence intensity was measured in one series of tissue for each DRG).

**Supplementary Figure 20.**
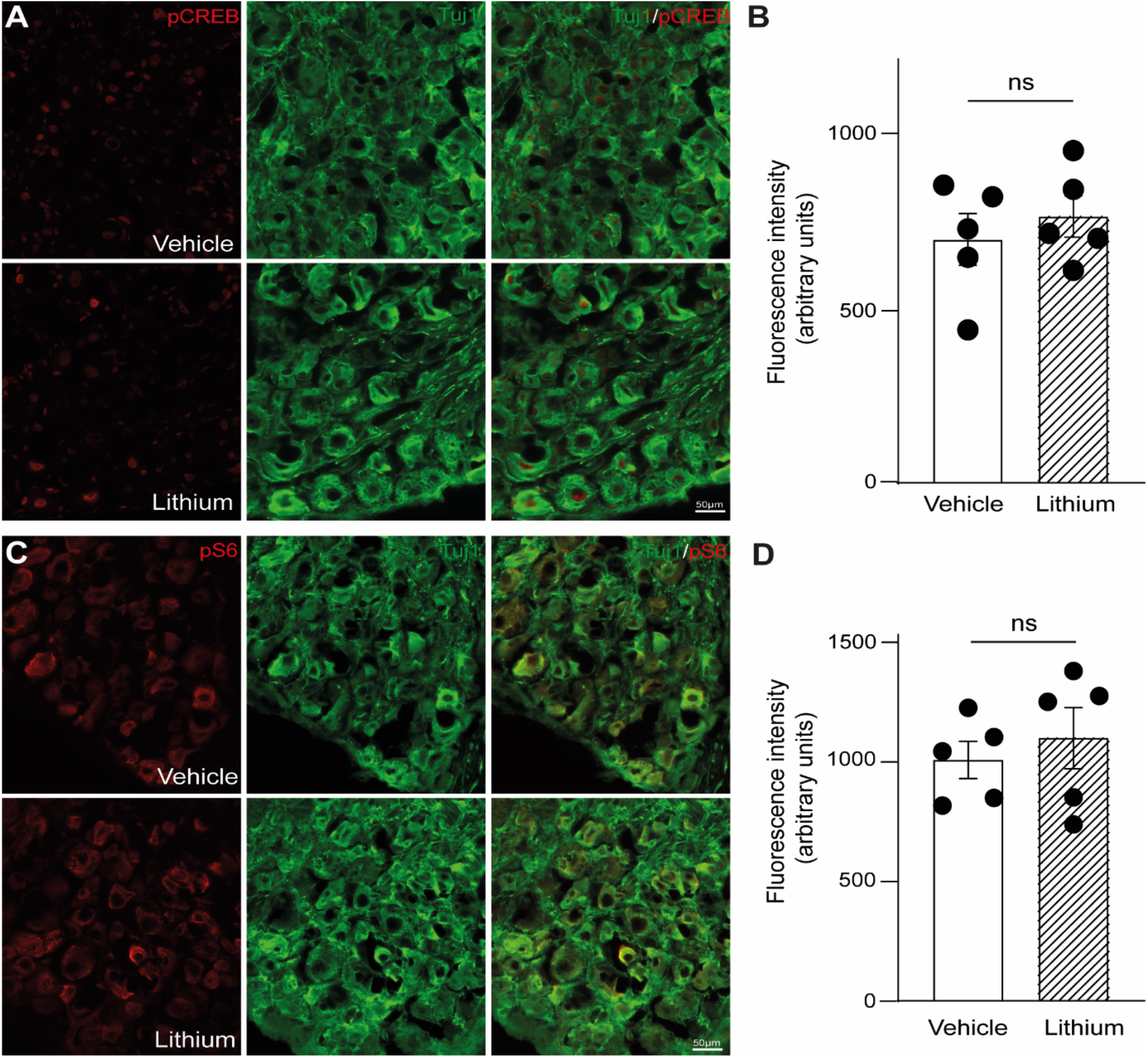
**A.** Representative images of DRG from WT mice treated with lithium or vehicle immunostained for pCREB (red) and Tuj1 (green). Scale bar, 50 µm. **B.** Quantification of the fluorescence intensity of pCREB (red) in Tuj1 positive DRG neurons (mean ± SEM, Student T-test, p<0.05, n=5 biologically independent animals/group). Fluorescence intensity was measured in one series of tissue for each DRG. **C**. Representative images of DRG from WT mice treated with lithium or vehicle immunostained for pS6 (red) and Tuj1 (green). Scale bar, 50 µm. **D.** Quantification of the fluorescence intensity of pS6 (red) in Tuj1 positive DRG neurons (mean ± SEM, Student T-test, p<0.05, n=5 biologically independent animals/group). Fluorescence intensity was measured in one series of tissue for each DRG.

## Methods

### Animals and Animal Husbandry

All animal experiments were carried out according to UK Home Office regulations in line with the Animals (Scientific Procedures) Act of 1986 under personal and project licenses registered with the UK Home Office. Mice were maintained in-house under standard housing conditions (12-h light/dark cycles). All mice were on a C57BL6 background (male, 20-30 g, 6-8 weeks of age). Bmal1^fl/fl^ (B6.129S4(Cg)-Arntltm1Weit/J; https://www.jax.org/strain/007668) and Per2-Luc mice (B6.129S6-Per2tm1Jt/J; https://www.jax.org/strain/006852) were purchased from Jackson’s laboratory, Cry1/2^-/-^ mice were gently provided by Dr. Marco Brancaccio. Experimenters were blind to experimental groups during analysis. Sample sizes were chosen in accordance with power analysis and with similar previously published experiments. The animals were housed in a controlled light dark cycle and housed individually. The animals were kept in constant conditions the day before the lesion.

### Sciatic nerve crush (SNC) surgery

Mice were anesthetized with isoflurane (5% induction, 2% maintenance), shaved on the hind limbs and lower back, cleaned with iodine, and an ophthalmic solution was put on the eyes to prevent drying. An incision was made on the skin and the biceps femoris and the gluteus superficialis were opened by blunt dissection and the sciatic nerve was exposed using a surgical hook. Sciatic nerve crush was performed orthogonally for 20 seconds (45 seconds for reinnervation experiment) using a 5 mm surgery forceps (91150-20 Inox-Electronic). The crush was performed at approximately 20 mm distally from the L4-L6 sciatic DRG.

### Sciatic nerve regeneration

24 hours or 72 hours following the surgery, sciatic nerves were dissected and post-fixed in 4% PFA, incubated at 4°C for 1h and transferred into 30 % sucrose for at least 3 days. Subsequently, the tissue was embedded and frozen in Tissue-Tek OCT and maintained at - 80°C until cut into 11 µm sagittal sections. Tissue sections were immunostained for SCG10 (1:1000, Rabbit, Novus) a marker for regenerating axons. The crush side was identified by deformation of the nerve and disruption of axon coinciding with highest SCG-10 intensity. The SCG-10 intensity was measured in 500 µm intervals along the length of the nerve distal to sciatic nerve crush side. The intensity was normalised to the SCG-10 intensity before the crush side and plotted as fold-change. 4-6 sections per animal were analysed and imaged with a HWF1 - Zeiss Axio Observer with a Hamamatsu Flash 4.0 fast camera using 10x magnification. The Regeneration index (RI) was calculated as the distance from the crush site at which the SCG-10 mean intensity equals 50% of the SCG-10 mean intensity at the crush site.

### Neurotrophin ELISA

Levels of neurotrophins were analysed in homogenised L4-6 DRG samples. DRGs were lysed in RIPA buffer and diluted 1:6. 100 µl of lysate (corresponding to 80 mg of proteins) were analysed with the Multi-Neurotrophin Rapid Screening ELISA kit (Biosensis), according to manufacturer’s instructions. The absorbance at 450 nm of the samples, obtained using GLoMax plate reader (Promega), was interpolated against the standard curve to calculate the neurotrophin concentration, and referred to mg of proteins.

### RT-qPCR

All RT-qPCR were performed using SYBR Green I (2×) from Platinum SYBR Green qPCR SuperMix-UDG kit (Thermo Fisher Scientific). Reactions were performed in triplicates on a 96-well plate with a final reaction volume of 20 μL by using 7900HT Fast Real-Time PCR System (Applied Biosystems, Foster City, CA USA). The thermal cycle profile consisted of 45 cycles divided in 4 stages (5 min at 95 °C, 20 s at 95 °C, 20 s at 56 °C, 20 s at 72 °C) and a final dissociation stage. The threshold cycle (Ct) data after each run was exported from SDS v2.4. The Ct data derived from the RT-qPCR were analysed using the ΔΔCt method, normalised against the endogenous reference gene GAPDH and expressed as fold change relative to the time point with highest expression or to control condition.

### Virus and CTB tracer delivery to DRG neurons

To transduce DRG neurons *in vivo*, 2 µl of AAV8-Cre-GFP or AAV8-GFP (Duke Viral Vector Core https://sites.duke.edu/dvvc/) were injected into the sciatic nerve of Bmal1flox mice (https://www.jax.org/strain/007668) 4 weeks prior to sciatic injury or collection using a 10 µl Hamilton syringe and Hamilton needle (NDL small RN ga34/15mm/pst45o) (Hamilton). For retrograde CTB tracing of regenerating sciatic fibres, 1 µl of CTB (List Biological Laboratories) diluted in saline were injected in the sciatic nerve distal to the lesion site using a 10 µl Hamilton syringe and Hamilton needle (NDL small RN ga34/15mm/pst45o) (Hamilton), 3 days before sacrificing the mice.

### RNA-sequencing

DRG samples were collected from animals that had undergone nerve injury at ZT8 and ZT20. Sciatic DRGs were extracted 72 h after injury (surgeries were performed as described above) and placed into RNAlater solution. The DRGs were crushed with an RNase-free micro-pestle, and neuronal enrichment was performed using a 15% BSA cushion prepared in a falcon tube: dissociated cells were carefully pipetted on top and then spun down at 80 x g for 8 min before being resuspended in culture media containing 1x B27 and Penicillin/Streptomycin in F12:DMEM. RNA was then immediately extracted using an RNAeasy Kit (Qiagen) according to the manufacturer’s guidelines. Residual DNA contamination was removed by treating the spin column with 40 units of RNase-free DNase I (Qiagen) for 15 min at 23 °C before RNA elution. RNA concentrations and purity were verified for each sample following elution with the Agilent 2100 Bioanalyzer. RNA with integrity number (RIN) factors above 8.5 were used for library preparation. cDNA libraries for each sample were generated by the Imperial BRC Genomics Facility using the TruSeq Sample Preparation Kit A (Illumina) and were sequenced using Illumina HiSeq 4000 (PE 2 × 75 bp) sequencing. GO analysis was performed on differentially expressed genes using DAVID 2021 (http://david.abcc.ncifcrf.gov/) and the dendrogram (heat map), PCA and odds ration were created and analysed using R. Differentially expressed genes were selected using a threshold of FDR < 0.01 and |1.5| < FC (fold change) or no FC cutoff.

### Intramuscular injection

Fourteen days post sciatic nerve crush sensory axons that had re-innervated the muscle were traced by injecting 5 µl of 1% CTB (List Biological Laboratories) diluted in saline into the tibialis anterior and medial gastrocnemius muscles using a 10 µl Hamilton syringe and Hamilton needle (NDL small RN ga34/15mm/pst45o) (Hamilton). 4 days post-tracing, mice were terminally anaesthetised and transcardially perfused with 20 ml PBS (pH 7.4) (Sigma) followed by 20 ml 4% paraformaldehyde in PBS (Sigma).

### Neuronal enriched Dorsal Root Ganglia (DRG) culture

Glass coverslips were coated with 0.1 mg/ml PDL, washed and coated with mouse Laminin 2ug/ml (Millipore) for 1-2 hours each previous to the start of the experiment. Sciatic DRG from adult animals were dissected and collected in Hanks balanced salt solution (HBSS) on ice. The DRG were transferred into a digest solution (5mg/ml Dispase II (Sigma), 2.5 mg/ml Collagenase Type II (Worthington) and incubated in a 37 °C water bath for 45 min, which occasional mixing. Thereafter, the DRG were washed and manually dissociated with a 1ml pipette in media containing 10 % heat inactivated FBS (Invitrogen) and 1x B27 (Invitrogen) in F12:DMEM (Invitrogen). Pipetting was continued until DRG were fully dissociated and no clumps could be observed. Next, the cell suspension was spun down at 1000 rpm for 4 min and resuspended in culture media containing 1x B27 and Penicillin/Streptomycin in F12:DMEM. For neuronal enriched cultures, a 15% BSA cushion was prepared in a falcon tube, dissociated cells were carefully pipetted on top and then spun down at 80 x g for 8 min before being resuspended in culture media containing 1x B27 and Penicillin/Streptomycin in F12:DMEM. 3500 cells were plated on each coverslip (laminin and PDL coated) and maintained in a humidified culture chamber with 5% CO2 at 37 °C, 24 hours before fixed with 4% PFA and immunostained or analysed for luminesce.

For in vitro luciferase assay, neuronal enriched DRG culture were synchronized with 100 nM of Dexamethasone for 2 h. The hormone was washed out and cultures were harvested at different time points for subsequent analyses.

### Lithium treatment after injury

An injection of lithium chloride i.p. (1 mEq/Kg) was given immediately after injury, then lithium carbonate (600 mg/L) was given in drinking water for 3 days from the day of surgery until sacrifice.

### Per2-Luciferase Detection and siRNA transfection of DRG cultures

DRG form Per2-Luc mice (https://www.jax.org/strain/006852) were cultured as previously described. siRNA transfection was performed 12 hours after plating according to manufacturer instructions. 72 hours after transfection, DRG cultures were lysated in 200ul of GloLysis buffer and an equal amount of GloMax luciferin reagent was added to the solution. Luciferase luminescence was assessed 24 hours after plating, every 4 hours across a 24 hours period using GloMax® Explorer Multimode Microplate Reader (GM3500) plate reader.

### Immunohistochemistry

Immunohistochemistry on tissue sections was performed according to standard procedures. Tissue sections were rehydrated with PBS for 5 minutes before blocking and permeabilization for 1 hour with 10% normal goat serum (Abcam) containing 0.3% PBS-TritonX-100. The sections were then incubated with anti-SCG10 (1:1000, Rabbit, Novus), anti-SOX10 (1:1000, Rabbit, Abcam), anti-CD68 (1:1000, Rabbit, Abcam), anti-Ly6G (1:500, mouse, BioxCell, clone 1A8), anti-Bmal1 (1:500, Rabbit Invitrogen, PA5-118391), anti-Tuj1 (1:500, Mouse, Novus NB100-1612) anti-CD68 (1:200, Rabbit, Abcam, ab125212), anti-NH200 (1:1000, Mouse, NovusBio, NB500-416), anti-PGP9.5 (1:200, Rabbit, Proteintech, 14730-1-AP), anti-CTb (1:1000, Goat, List biological 703), anti-GFP (1:1000, Chicken, Abcam ab6556), H3K27ac (1:2000, Rabbit, Abcam, Abcam ab4729), anti-CGRP (mouse, abcam ab81887, 1:100), anti-pCREB (1:500, Rabbit, CST, #9198), anti-pS6 (1:1000, Rabbit, CSR, #35708) at room temperature over-night. The sections were washed three times with PBS, followed by incubation with Alexa Fluor conjugated goat secondary antibodies for 1 hour. All tissue sections were counterstained with DAPI (Molecular Probes) and cover slipped using mowiol.

### Immunocytochemistry (ICC)

Plated cells were fixed by incubation with cold 4 % PFA for 15 min. Thereafter, they were blocked and permeabilized for 1 hour with 0.3 % TX100 in PBS containing 2 % BSA. The primary antibody staining was performed using anti-Bmal1 (1:500, Rabbit Invitrogen, PA5-118391), anti-anti-βIII tubulin (1:1000, Mouse, Novus NB100-1612), anti-GFP (1:1000, Chicken, Abcam ab6556) in 0.1 % TX100 in PBS containing 2 % BSA, which O/N incubation at RT. The goat secondary antibody (Alexa) was diluted in 0.1 % TX100 in PBS containing 2 % BSA and cells were incubated for 1 hour. All cells were counterstained with DAPI.

### Microscopy

Photomicrographs were taken with a Nikon Eclipse TE2000 microscope with an optiMOS scMOS camera using 10x or 20x magnification - Zeiss Axio Observer with a Hamamatsu Flash 4.0 fast camera using 10x or 20x magnification. Confocal images were taken with a Leica TCS SP8 II confocal microscope at 20X magnification and processed with the LAS-AM Leica software (Leica).

### Image Analysis for IHC and ICC

Image analysis was conducted using ImageJ (Fiji) software. All analysis was performed in blind to the experimental groups.

DRG images were taken using a Nikon Eclipse TE2000 microscope with an optiMOS scMOS camera at either 10x or 20x magnification. Images were analysed by calculating the percentage of cells with positive staining.

For neurite length analysis between 15 and 20 images were taken per coverslip and analysed using NeuronJ plugin for Image J software (Image J). All analyses were performed in blind. Approximately 45-60 cells were analysed per animal and condition.

### Statistical analysis

Results are charted as mean ± SEM. Statistical analysis was performed using GraphPad Prism 8. Normally distributed data were evaluated using a two-tailed unpaired Student’s t-test or a one-way ANOVA when experiments contained more than two groups. Tukey’s or Sidak’s test or multiple comparison testing corrected by FDR with Benjamini and Hochberg were applied when appropriate. The two-way ANOVA, Tukey’s or Sidak’s test, was applied when two independent variables on one dependent variable were assessed. A threshold level of significance was set at P<0.05. Significance levels were defined as follows: * P<0.05; ** P<0.01; *** P<0.001; **** p<0.0001. All data analysis was performed blind to the experimental group.

Rhythmicity of RT-PCR data (Figure 1, p<0.05) and Per2 Luciferase oscillation upon Bmal1 deletion in vitro (Figure 2, p<0.005) was assessed in Biodare2^43^ by using JTK_CYCLE^44^ on mean values, with dataset compared to a 24h cosine wave with a 4h phase spread.

